# Freshwater Microbiomes Shape Viral Inactivation in Continuous Cultures

**DOI:** 10.64898/2026.09.26.754611

**Authors:** L. Daniela Morales, Joe Heffron, Rachel M. Cook, Htet Kyi Wynn, Mark A. Borchardt, Hannes Peter, Tamar Kohn

## Abstract

In freshwater environments, bacteria are known to enhance the removal and inactivation of human viruses. Rates of inactivation differ across freshwater ecosystems, though the influence of community diversity and composition on this process remain poorly understood. Here, we used dilution-to-extinction to generate 64 distinct freshwater communities from three lakes and three aquifers. After establishing freshwater communities in chemostat cultures, each community was challenged with two viruses, coxsackievirus B5 and human adenovirus 2. Both viruses were inactivated more rapidly in lake-derived than in groundwater-derived communities, and adenovirus was inactivated faster than coxsackievirus in both freshwater types. Community-level parameters (e.g., cell numbers, richness) were not significant predictors of viral inactivation. Instead, differential abundance analyses identified bacterial OTUs associated with inactivation: *Acidovorax* sp. were associated across sources and *Brevundimonas* sp., *Sphingopyxis* sp., and *Hydrogenophaga* sp. showed source-specific associations. Together, these findings provide new insights into the ecological drivers of viral stability and suggest that viral inactivation may depend more strongly on specific microbial taxa and their environmental context than on overall community diversity. Identifying these associations improves our understanding of the factors governing viral stability in freshwater environments.

**GRAPHICAL ABSTRACT:** 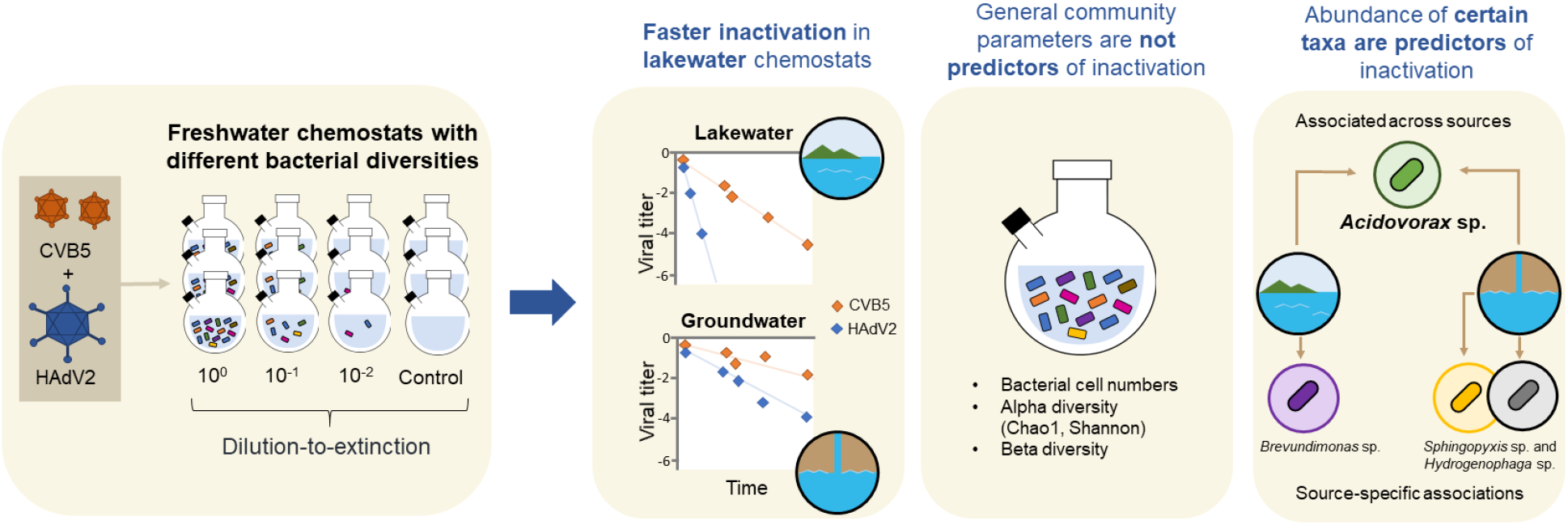

## INTRODUCTION

Enteric viruses withstand conventional wastewater treatment barriers and contaminate aquatic environments, where they can persist and seed infectious outbreaks ^1–3^. Their stability in the environment is challenged by both abiotic (e.g., temperature, sunlight) and biotic (e.g., microorganisms, extracellular enzymes) factors. Nevertheless, many viruses remain infectious for weeks to months under favorable conditions, presenting a sustained public health threat ^4,5^.

Several studies show that microorganisms play an important role in viral inactivation in freshwater ecosystems ^6–8^. However, the underlying metabolic and ecological processes that drive inactivation remain incompletely understood. Most mechanistic studies to date have focused on individual bacterial isolates or specific antiviral compounds^7,9,10^. For example, Corre *et al*. (2022), found that bacterial cells are required for the inactivation of echovirus 11 and coxsackievirus A9 in lakewater and that several bacterial isolates from Lake Geneva exhibited antiviral activity against both viruses. The extent of the observed inactivation depended on both the viral and the bacterial species^11^. In contrast, the role of complex microbial communities in regulating virus inactivation is less explored. Interaction among bacterial taxa can influence community structure, metabolic activity, and the production of extracellular compounds, thereby potentially enhancing or reducing viral inactivation. Whether and how microbial community composition and diversity shape viral inactivation in freshwater ecosystems remains largely unexplored.

Microbial diversity is a critical determinant of ecosystem functioning. While some ecosystem functions are shared across taxa (e.g., decomposition of organic matter, aerobic respiration and fermentation), others are taxa-specific (e.g., methanogenesis, nitrogen fixation)^12^. Identifying microbial communities or taxa associated with a specific function is essential to understand the mechanisms that drive these processes. We have previously explored viral inactivation as an ecosystem function in lakewater^13^. Using batch cultures from Lake Geneva, we showed that the capacity to inactivate echovirus 11 was weakly associated with bacterial community richness and abundance of certain taxa. However, whether this function is widely spread across freshwater ecosystems remains unknown. Comparing freshwater communities from different sources will therefore provide an opportunity to determine whether viral inactivation is a general property of freshwater microbiomes and to identify bacterial taxa associated with this function.

Here, we aimed to compare the viral inactivation capacity of groundwater and lakewater communities and to investigate the role of bacterial diversity and composition on viral inactivation. To achieve this, we collected water from three Swiss lakes and three aquifers in Wisconsin, USA, and used a dilution-to-extinction approach to generate 64 microbial communities with diversity gradients in chemostat cultures. The communities were challenged with two viruses (coxsackievirus B5 (CVB5) and Human Adenovirus 2 (HAdV2)) to assess their viral inactivation capacity. Viral inactivation was analyzed as a function of microbial diversity estimates and community composition. Our results further characterize viral inactivation as an ecosystem function and identify candidate bacterial biomarkers of this process in freshwater, including biomarkers shared across groundwater and lakewater, and specific to each source.

## MATERIALS AND METHODS

### Sampling of lakewater and groundwater

Lakewater samples were taken from three Swiss lakes (**Figure 1A**): Lake Geneva was sampled on February of 2025 (46.51336° N, 6.57219° E); Lake Bret on October of 2025 (46.50946° N, 6.76862° E); and Lake Brienz on December of 2025 (46.68867° N, 7.90737° E). Around 30 L of water were taken from the surface of each lake approximately 3-5 m from the shore and stored in previously autoclaved carboys. Carboys were rinsed twice with lakewater prior to collection. Conductivity, pH, dissolved oxygen and temperature were measured directly on each site using a digital meter Multi 3430 (WTW) (**Table S1**) before sample collection. Lakewater was immediately transported to the laboratory and stored at 4°C until experimental use. A sample from each lake was taken to determine total organic carbon (TOC) and total nitrogen (TN) content using a Vario TOC Cube (Elementar) (**Table S1**).

**Figure 1.**
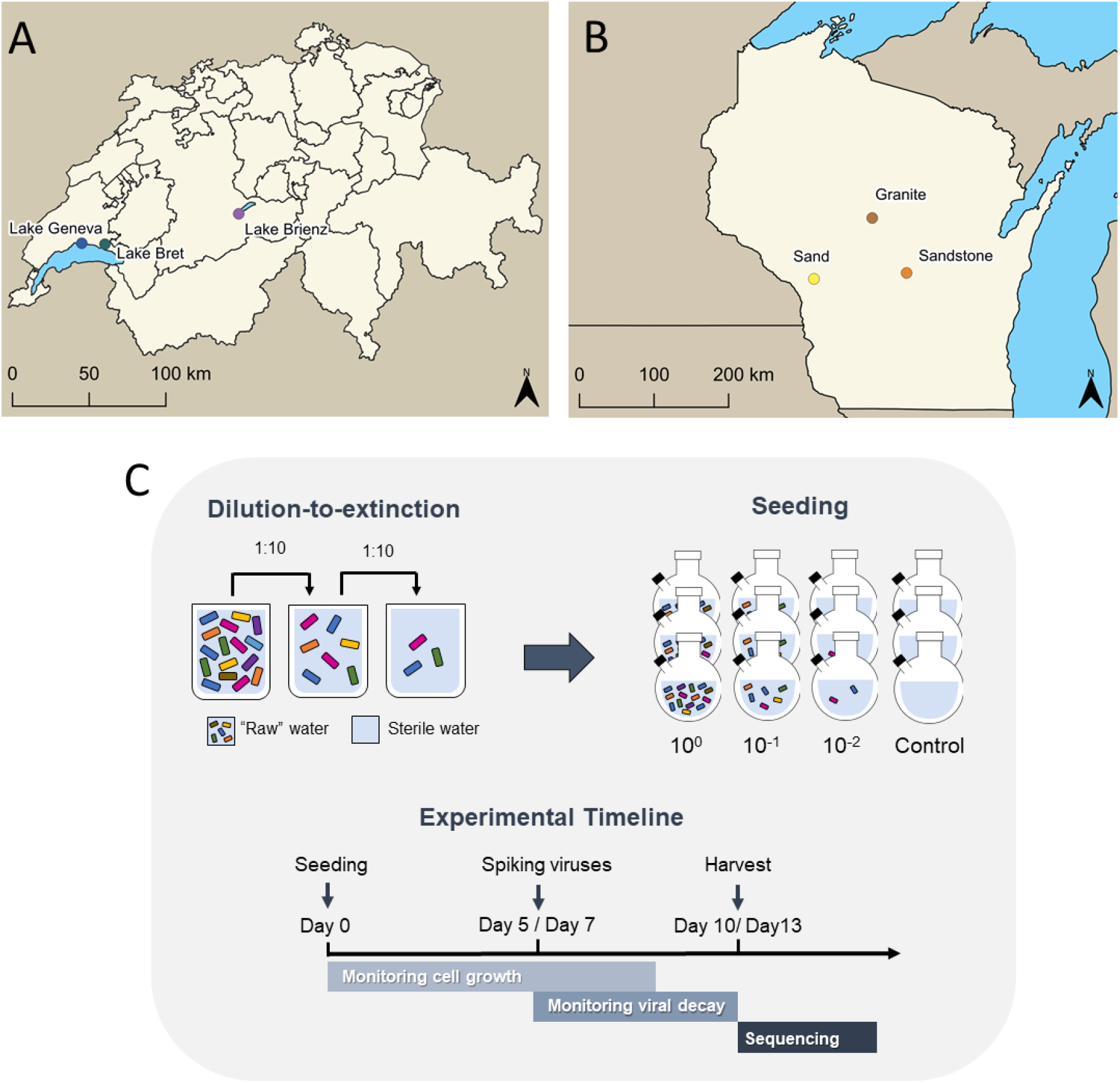
Water sampling and experimental set up. Lakewater **(A)** samples were taken from three Swiss lakes: Lake Geneva, Lake Bret and Lake Brienz. Groundwater **(B)** samples were taken from three aquifers with different geology located in Melrose (Sand), Hancock (Sandstone) and Stratford (Granite) (Wisconsin, USA). **(C)** Chemostats were seeded with undiluted (10^0^) or serially diluted (10^-1^, 10^-2^) lakewater or groundwater. Once communities were established (5 days for lakewater; 7 days for groundwater), a mixture of Coxsackievirus B5 (CVB5) and Human Adenovirus 2 (HAdV2) was spiked into each chemostats, effluent was sampled over time and viral infectivity was monitored. Unseeded controls were maintained alongside seeded chemostats. To assess community composition, 16S ribosomal RNA gene was amplified and sequenced.

Groundwater samples were taken from three aquifers located in Wisconsin (USA) (**Figure 1B**): a granite aquifer (44.76° N, 90.10° W; sampled May 2025), a sand aquifer (44.05° N, 91.05° W; sampled December 2025), and a sandstone aquifer (44.12° N, 89.54° W; sampled December 2025). Aquifers were sampled from flame-sterilized taps downstream of the pressure tank but upstream of any treatment or conditioning. Aquifer water was collected in autoclaved 20 L carboys after flushing a minimum of 40 L. Groundwater was immediately transported to the laboratory and stored at 4°C until experimental set up. Conductivity and pH were measured upon return to the laboratory using a Traceable Expanded Range Conductivity Meter (Fisherbrand) and Orion Star A211 (Thermo Scientific), respectively (**Table S1**). A sample from each well was taken to determine total organic carbon (TOC) and total nitrogen (TN) using a TOC analyzer (Teledyne Tekmar Torch, Mason, OH) and an automated flow analyzer (Quikchem 8500 [method 10-107-04-1-C]; Hach, Loveland, CO.) (**Table S1**).

Before each experiment, an aliquot of each source water was prepared for 16S rRNA sequencing (lakewater: 200 mL; groundwater: 2 L). Lakewater samples were vacuum filtered using 0.22 μm polycarbonate track-etched filters (Sartorius) and stored at −20°C for DNA extractions. Groundwater samples were amended with Tween 20 (Thermo Scientific, Fair Lawn, NJ, USA) to a final concentration of 0.1% (v/v) and filtered with 0.2 μm CP Select concentrating pipettes (InnovaPrep, Drexel, MO, USA). Concentrating pipets were eluted with Tris Elution Buffer (InnovaPrep) and stored at −80°C until analysis. In addition, water from each source was sterilized by ultrafiltration and ultraviolet (UV) radiation using a dead-end Rexeed-25A hemodialyzer (Asahi Kasei Medical) and a UV treatment unit (Viqua model VH150, Guelph, Ontario; flow rate of 1 L min^-1^, yielding a dose of approximately 520 mJ/cm^2^). This water was used to feed chemostat cultures and dilute starting communities.

### Chemostat cultures

To generate bacterial communities of different diversities we used a dilution-to-extinction approach. This method reduces total abundance in a community in successive dilutions, which tends to select against rare species while retaining only initially abundant species in more dilute samples ^14^. For this approach, lakewater or groundwater from each source was serially diluted into sterile water from the same source. Replicate chemostats (*n* = 2-3) were seeded with 400 mL of a given dilution (10^0^, 10^-1^, 10^-2^) and grown for 10 to 14 days using sterile source water to feed the cultures (**Figure 1C**). As a control, every run included chemostats filled with sterile source water only. Control chemostats were spiked with antibiotics (1% Penicillin-Streptomycin (GibcoTM) or 500 ug mL^-1^ of cefotaxime + 100 ug mL^-1^ of ampicillin) every other day during the experiments to avoid bacterial growth. Lakewater chemostats were fed at a rate of 0.125 mL min^-1^ and groundwater chemostats at 0.075 mL min^-1^, consistent with bacterial growth rates encountered in lakewater and groundwater ^15–17^. Resulting dilution rates in chemostats were 0.02 h^-1^ and 0.01 h^-1^ in lakewater and groundwater chemostats, respectively. We previously found these rates to yield high microbial diversity while minimizing washout of virus^14^. The effluent pump was set to a speed 20% higher than that of the influent pump (0.150 mL min^-1^ and 0.09 mL min^-1^) to avoid overflowing. Chemostats were grown aerobically using filtered air and incubated in the dark at room temperature (22°C). Effluent volumes were measured daily to ensure proper function of pumps at the desired rate. On the last day of the run, the content of each chemostat was filtered and preserved for 16S rRNA sequencing following the same procedure as source water samples.

### Monitoring bacterial growth in chemostats

To monitor bacterial growth inside chemostat cultures, 0.9 mL of the effluent was sampled every two to three days, fixed with 0.1 mL of 37% filtered formaldehyde (0.22 μm) and stored at 4°C until analysis. For lakewater, the number of bacteria was quantified using a volumetric flow cytometer Attune CytPix (ThermoFisher Scientific, Waltham, MS, USA) or a NovoCyte (Acea). For this, 198 μL of the fixed sample was stained with 2 μL of 20 μM Syto13 nucleic acid stain (ThermoFisher Scientific) followed by incubation in the dark at room temperature for 15 min. Groundwater samples were analyzed using a CytoFLEX V2-B2-R0 (Beckman Coulter) stained with SYTO16 (10 μM). Bacterial cells were identified using fluorescence and side scatter properties. Finally, cell counts were adjusted accounting for fixative and nucleic acid stain used during sample preparation.

### Virus propagation and enumeration

Coxsackievirus B5 (CVB5) was propagated from an environmental isolate (GenBank accession number MG845891) and Human Adenovirus 2 (HAdV2) was kindly provided by Rosina Girones (University of Barcelona, Spain). HAdV2, a respiratory type, was used as a model for enteric adenoviruses such as Human Adenovirus 40 and 41. CVB5 and HAdV2 viral stocks were prepared in Buffalo Monkey Kidney cells (BGMK). For viral propagation, cells were grown at 37°C and 5% CO_2_ with Minimum Essential Medium (MEM, A41922) supplemented with 1% penicillin-streptomycin (15140, Gibco) and 10% heat-inactivated fetal bovine serum (FBS, A5256701, Gibco). Once cells reached 90% confluence, growth media was discarded and cells were maintained in MEM supplemented with 1% penicillin-streptomycin (15140, Gibco) and 2% heat-inactivated fetal bovine serum (FBS, A5256701, Gibco). A viral suspension of either CVB5 or HAdV2 was added to the cells and incubated for 3-5 days. Next, viruses were released from the cells by freeze-thawing the culture flasks three times and cell debris were removed by centrifugation at 1’100x g for 10 minutes. Virus stocks were prepared by buffer-exchange into phosphate-buffered saline (PBS, 18912014, Gibco) using Amicon Ultra-15 centrifugal filters (UFC910024, Merck Millipore).

Infectious viruses were enumerated as previously described by the Most Probable Number method^18^. BGMK cells were used to assess the infectivity of both viruses combined, whereas Rhabdomyosarcoma (RD) cells were used for HAdV2 only. This approach was possible because RD cells were only infected by HAdV2, but not by the environmental isolate of CVB5 used herein. Briefly, each cell line was grown in 96-well plates (Greiner CELLSTARVR 96-well plates; Sigma Aldrich) until 90-95% confluent and five replicates of each sample were then 10-fold serially diluted into the cells. The plates were incubated at 37°C in 5% CO_2_ for 5 days. Each well was examined under the microscope for cytopathic effect (CPE). The number of CPE-positive wells was converted to most probable number per mL (MPN/mL) using the R package MPN. Samples were considered as quantifiable if at least one of the five wells containing the lowest dilution of a given sample yielded a positive CPE. The limit of quantification (LOQ) corresponded to one positive well (of five replicates) at the highest dilution (10-fold diluted) and corresponded to 90.4 MPN/mL.

### Virus inoculation

Once bacterial communities reached stable cell densities inside the chemostat cultures (5 days for lakewater; 6-7 days for groundwater^14^), 4 mL of a viral suspension were spiked into each chemostat to a final concentration of 10^6^ MPN/ml of CVB5 and HAdV2. After allowing chemostats to mix for 1 or 2 hours, samples (1-2 mL) were taken from the effluent to estimate viral infectivity over time for up to 10 days for lakewater chemostats and 13 days for groundwater chemostat, and stored at −20°C until enumeration.

### Kinetic data analysis

Viral inactivation rate constants were determined by fitting a first-order decay model to the decay curve (Equation 1):

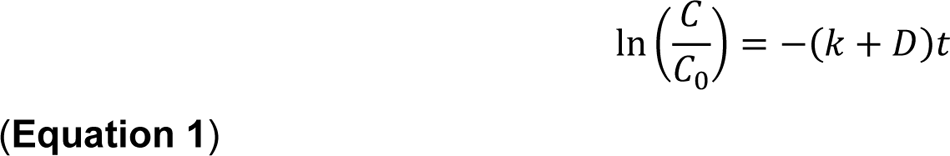

where *C* (MPN/mL) are infectious virus concentrations at time *t* (h) after the virus spike and *C_0_* (MPN/mL) are infectious virus concentrations at the first measured time point in chemostats, *k* is the inactivation rate constant (h^−1^), and *D* is the dilution rate (h^-1^). Because HAdV2 was generally more rapidly inactivated in all chemostats and often was spiked at a slightly lower concentrations compared to CVB5, inactivation kinetics obtained in BGMK cells were attributed to CVB5, while data obtained in RD cells were attributed to HAdV2. However, as HAdV2 may also be detected in BGMK cells, we cannot exclude the possibility that the infectivity measurements attributed to CVB5 were partially influenced by HAdV2 in some chemostats.

For rapidly inactivating chemostat, the virus titer could fall close to or below the LOQ already at the first sampling time point. Thus, a minimum rate constant (*k*_min_) was determined according to

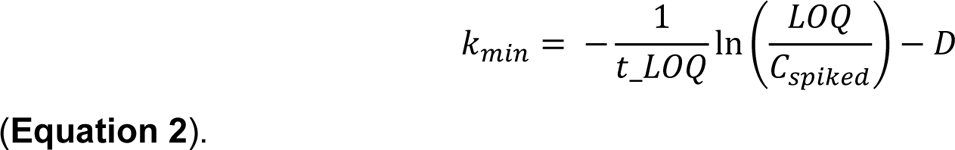

where LOQ is the limit of quantification (MPN/mL), *C_spiked_* is the spiked virus concentration, *t_LOQ* (h) is the time at which the virus concentration becomes undetectable, and *D* is the dilution rate (h^-1^).

### 16S rRNA gene amplicon sequencing and microbial diversity analyses

Total DNA was extracted from samples using the DNeasy PowerWater Kit (Qiagen, Hilden, Germany) according to the manufacturer’s instructions. Extraction blanks (extraction following the manufacturer’s protocol without any input material) were included in every run. DNA extracts were quantified using Qubit and the dsDNA HS Assay (ThermoFisher Scientific) and checked for purity using microvolume spectrophotometer (NanoDrop, ThermoFisher Scientific or N120 Nanophotometer, Implen).

DNA extracts were then used for 16S rRNA gene amplification. The PCR reaction was carried out in 30-50μL reactions using LongAmp Hot Start 2X Master Mix (New England Biolabs, Ipswich, MS, USA) and 2 μL of 27F (5′-AGAGTTTGATCMTGGCTCAG-3′) and 1492R (5′-CGGTTACCTTGTTACGACTT-3′) primer pair at a concentration of 10 μM. The PCR conditions were as follows: initial denaturation at 94 °C for 1-3 min, 30 cycles of 94 °C for 30 s, 55 °C for 30-45 s, 65-68 °C for 1-2 min with a final extension at 65-68 °C for 5 min. Subsequently, each PCR product was purified using Agencourt AMPure XP beads (Beckman Coulter, Brea, CA, USA) and quantified using Qubit (dsDNA quantification, Broad Range) or an N120 Nanophotometer (Implen). The sequencing library was prepared using the Native Barcoding Kit SQK-NBD114.24 (Oxford Nanopore Technologies, Oxford, UK) following the manufacturers’ instructions. The library was loaded into the FLO-MIN114 flow cell (Oxford Nanopore Technologies). Flow cells were run on a MinION Mk1C or Mk1B sequencing device for 72 h or the life of the flow cell. Further details on sequencing data processing are included in supporting information **(Supplemental Method S1)**.

### Identification of biomarkers for virus inactivation

To identify differentially abundant bacterial species associated with viral inactivation, Analysis of Compositions of Microbiome with Bias Correction (ANCOM-BC) was performed using the *phyloseq* (version 1.52.0) and *ANCOMBC* (version 2.13.1) packages in R^19,20^. OTU-specific log-fold changes were estimated using regression models with calculated rate constants (*k)* included as continuous predictors. Analyses were conducted separately for lakewater and groundwater. For lakewater, pseudo-count sensitivity was used to assess the robustness of associations, whereas this criterion was not applied in the groundwater analysis due to the smaller gradients of inactivation. To account for multiple comparisons across taxa, *p* values were adjusted using the Benjamini–Hochberg false discovery rate (FDR) procedure. Taxa with an FDR-adjusted q-value < 0.05 were considered significantly associated with *k*. Positive regression coefficients (β > 0) indicated that taxon abundance increased with increasing *k* values, whereas negative coefficients (β < 0) indicated that taxon abundance decreased with increasing *k* values.

### Identification of predictors of viral inactivation using multiple linear regression models

In addition to the abundance of candidate biomarkers identified using ANCOM-BC, we determined if high-level community properties (e.g., diversity indices, composition, certain taxa abundance) explained virus inactivation. Multiple linear regression models were fitted separately for groundwater and lakewater chemostats. The response variable was *k* and predictor variables included bacterial cell abundance, bacterial diversity indices (Chao1 and Shannon), microbial community composition (PCo1 and PCo2), and the relative abundance of selected bacterial taxa that were part of the top 20 on ANCOM-BC analyses. Prior to model fitting, pairwise Pearson correlations among all candidate predictors were assessed to identify collinearity. Where two predictors were strongly correlated (correlation coefficient > 0.7), only one was retained in the model. For taxa-level predictors, OTUs assigned to the same genus and showing a consistent (positive) association by ANCOM-BC analyses were combined into a single relative-abundance variable prior to correlation screening, as their individual abundances were highly redundant. The final set of non-redundant predictors was then used to fit the model for each response variable. Multiple linear regression models were fitted using the *lm()* function in R. Standardized regression coefficients (β) were calculated to allow comparison of the relative strength of associations among predictors measured on different scales. Model coefficients were visualized with their 95% confidence intervals. Predictors with positive standardized coefficients were interpreted as being positively associated with viral inactivation, whereas negative coefficients indicated an inverse association. Statistical analyses were performed in R.

## RESULTS

### Environmental and microbial parameters across source waters

We cultured microbial communities obtained from six different source waters: three lakes and three aquifers (**Figure 1A, B**). The environmental parameters measured in each source water at the time of sampling are shown in **Table S1**. We chose three lakes with different trophic states, sizes and locations: Lake Geneva and Bret are classified as mesotrophic lakes while Brienz is oligotrophic ^21^. For source aquifers, we selected wells representing aquifers with different geology (granite, sandstone and sand).

Using 16S rRNA gene sequencing, we detected a total of 21025 OTUs: 9602 (41.1%) were specific to groundwater and 10926 (51.6%) to lakewater. Only 497 (7.4%) OTUs were shared between them (**Figure S1A**). Alpha diversity analyses showed greater richness in lakewater (Chao1 = 5782 ± 1058) compared to groundwater (Chao1 = 3860 ± 842) (**Table S2**). Analysis of Bray-Curtis distance matrices using Principal Coordinate Analyses (PCoA) showed clear separation between lakewater and groundwater along PCo1 (24% variance) (**Figure S1B**). Groundwater samples exhibited greater within-group variation and were more dispersed along PCo2 (20.2% variance) compared to lakewater samples. In addition, we observed differences in relative abundances across source waters. *Limnohabitans* sp. were mostly abundant across lakewater samples, while *Aquabacterium* sp. and *Denitratisoma* sp. were predominant in groundwater samples (**Figure S1C**).

### Dilution-to-extinction generated freshwater communities with different community diversity and compositions

We used a dilution-to-extinction approach to generate communities with different microbial diversities and compositions (**Figure 1C**). Although initial cell densities differed among seeded dilutions, cell densities reached similar levels within 5-7 days and remained stable throughout the experiments (**Figure S2 and S3**). After regrowth, we observed higher cell densities in lakewater chemostats compared to groundwater. Lakewater chemostats reached cell densities between 10^5^ – 10^7^ cells mL^-1^, and groundwater chemostats grew to densities between 10^4^ - 10^5^ cells mL^-1^. Unfortunately, despite the use of different antibiotic combinations and doses, we observed bacterial growth in our unseeded controls.

At the end of each chemostat run, we performed 16S rRNA sequencing of all chemostat cultures (n = 64). As antibiotic treatment of unseeded controls resulted in communities that differed markedly from those of the seeded cultures (**Figure S4**), we excluded these chemostats from community comparison analyses. PCoA based on Bray-Curtis distance matrix of experimental chemostat cultures showed the greatest variation between lakewater and groundwater chemostats (19.6% variance) (**Figure 2A**). Analyses of lakewater and groundwater chemostats independently showed that chemostat communities clustered according to the source waters, and that dilution-to-extinction generated compositional shifts. Mean pairwise Bray-Curtis dissimilarities between different source waters were high for both groundwater (0.94 ± 0.04) and lakewater (0.86 ± 0.04) (**Table S3**). In contrast, dissimilarities were generally lower when comparing communities from the same source across different starting dilutions, with mean values ranging from 0.71 to 0.80 for groundwater and from 0.67 to 0.84 for lakewater chemostats (**Table S4**). A noted exception was observed in chemostats from Lake Brienz which exhibited pronounced compositional shifts across dilutions (**Figure 2A**). Bray–Curtis dissimilarities between the undiluted and 10⁻¹ and 10⁻² dilution communities were 0.89 ± 0.04 and 0.92 ± 0.03, respectively, whereas the two diluted communities were more similar to each other (0.72 ± 0.03).

**Figure 2.**
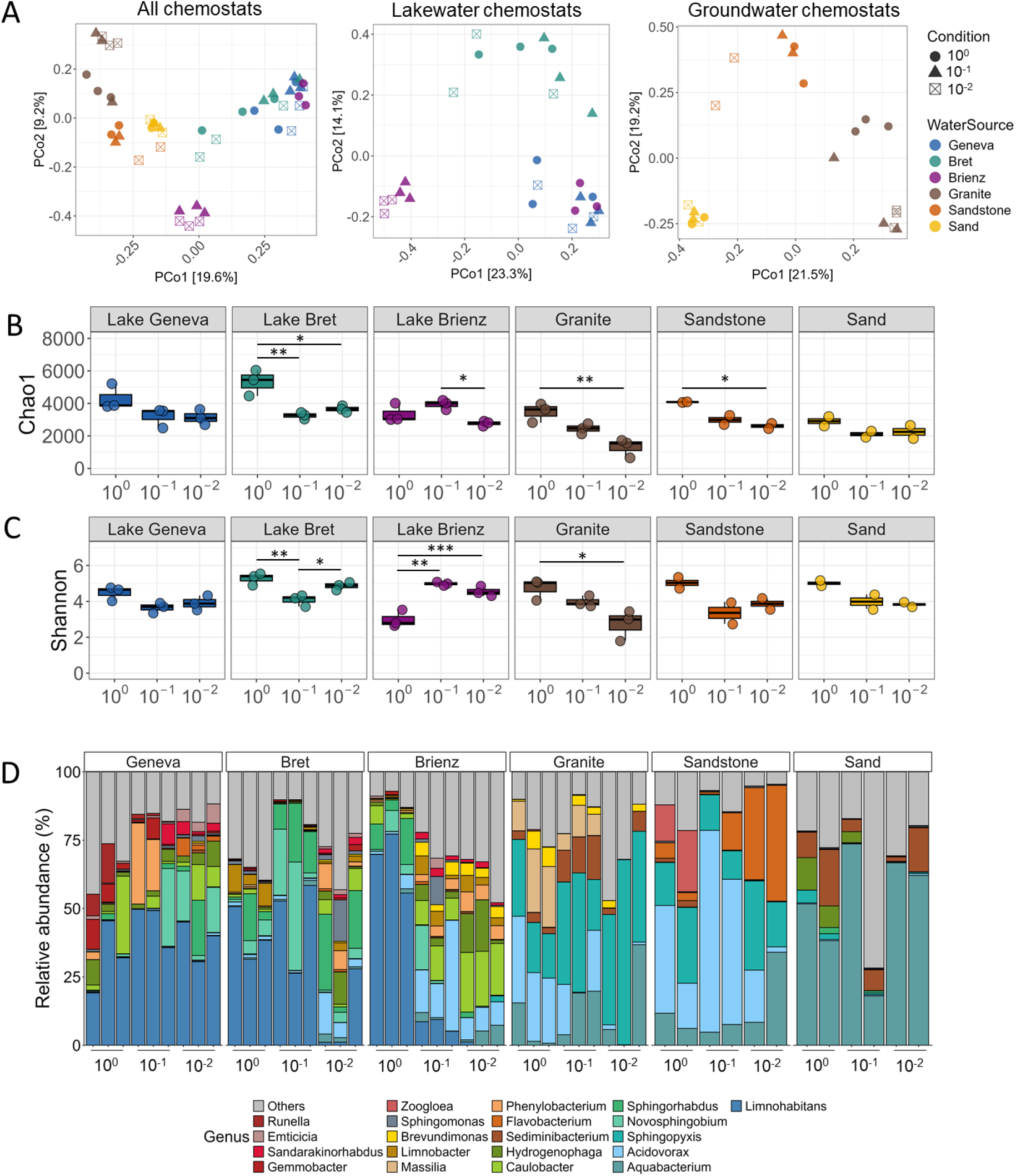
Microbial diversity comparison of chemostat communities. (A) Principal coordinate analysis (PCoA) of Bray–Curtis distances calculated from all chemostat communities, comparing bacterial community composition across all cultures and separately among groundwater and lakewater cultures. Calculated Chao1 (B) and Shannon (C) indexes per chemostat, averaged per culture starting dilution and water source. (D) Relative abundance of the top 20 most abundant bacterial genera grown inside chemostat cultures. Chao1 and Shannon indices were compared among conditions within each water source using one-way ANOVA. Significance codes: 0 ‘***’ 0.001 ‘**’ 0.01 ‘*’ 0.05.

Alpha diversity analysis showed that dilution decreased both Chao1 (estimating total richness) and Shannon indices (incorporating community richness and evenness) (**Figure 2B-C**). Chao1 declined significantly with increasing dilution in lakewater chemostats seeded from Lake Bret and in groundwater chemostats seeded from granite and sandstone aquifers (p < 0.05 for all comparisons; **Table S5**). Lake Brienz chemostats showed the opposite pattern for Shannon diversity, with 10⁻¹ and 10⁻² dilutions exhibiting significantly greater values than undiluted (10⁰) cultures (p < 0.01). Beyond the effects of dilution, alpha diversity was generally lower under chemostat conditions than in the source communities, with reductions in Chao1 of 6–45% and in Shannon diversity of 4–58% (**Tables S2, S5–S8**), with a few exceptions—chemostats from undiluted Lake Bret, sand, and sandstone water, maintained richness similar to, or higher than, their source water communities.

We next qualitatively examined community composition across chemostats (**Figure 2D**). *Limnohabitans sp.* and *Aquabacterium sp.* remained the most prevalent taxa in lake- and groundwater chemostats, respectively, and *Acidovorax* sp. was consistently among the dominant taxa across most chemostats under our selected growth conditions, with the exception of chemostats from the sand aquifer and Lake Geneva. Dilution-to-extinction was associated with source-water-dependent shifts in the dominant community members: in Lake Bret and Brienz chemostats, *Limnohabitans* sp. appeared less prevalent with increasing dilution, while *Acidovorax* sp., *Caulobacter* sp., and *Hydrogenophaga* sp. became more prevalent. These qualitative compositional shifts are consistent with the reduced Chao1 and increased Shannon values previously observed for Lake Brienz chemostats (**Figure 2B-C**). Groundwater chemostats showed comparatively less variation in dominant taxa across dilutions, with *Aquabacterium* sp. prevalent in most chemostats. In addition, *Acidovorax* sp. appeared less prevalent in chemostats seeded from diluted granite and sandstone aquifers. Overall, comparisons of alpha and beta diversity, as well as qualitative assessment of community composition, indicate that source water had a greater influence on community composition than seeding dilution, and that dilution-to-extinction altered the relative prevalence of dominant taxa while community signatures characteristic of each source were largely maintained.

### Lakewater chemostats showed faster viral inactivation compared to groundwater

Once chemostat communities reached a constant cell density (5 days for lakewater chemostats and 6-7 days for groundwater; see **Figure 3A** for average cell density in chemostat cultures measured within one day of virus inoculation), a mixture of CVB5 and HAdV2 was directly spiked into the chemostats. We observed faster inactivation of HAdV2 compared to CVB5 in all chemostats, and faster inactivation in chemostats seeded from lakewater compared to those from groundwater (**Figure S5, S6, S7 and S8**). In lakewater chemostats, the inactivation rate constants for CVB5 (*k*_CVB5_) ranged from 0.11 to 0.22 h^-1^ with the fastest inactivation found in chemostats from Lake Brienz (**Figure 3B; Table S9**). As most of HAdV2 was inactivated below the LOQ in lakewater chemostats within the first 4 hours (**Figure S7**), we estimated the minimum rate constant (*k_min HAdV2_*) instead of *k*_HAdV2_ (**Figure 3C**), which fell in the range of 0.54 – 7.12 h^-1^. Similar to our observations for *k*_CVB5_, chemostats from lakes Brienz and Bret had faster HAdV2 inactivation compared to Lake Geneva (**Figure S7**). In groundwater chemostats, *k*_CVB5_ ranged from 0 to 0.06 h^-1^ and *k*_HAdV2_ from 0.01 to 0.93 for h^-1^ (**Figure 3B and 3D; Table S10 and S12**).

**Figure 3.**
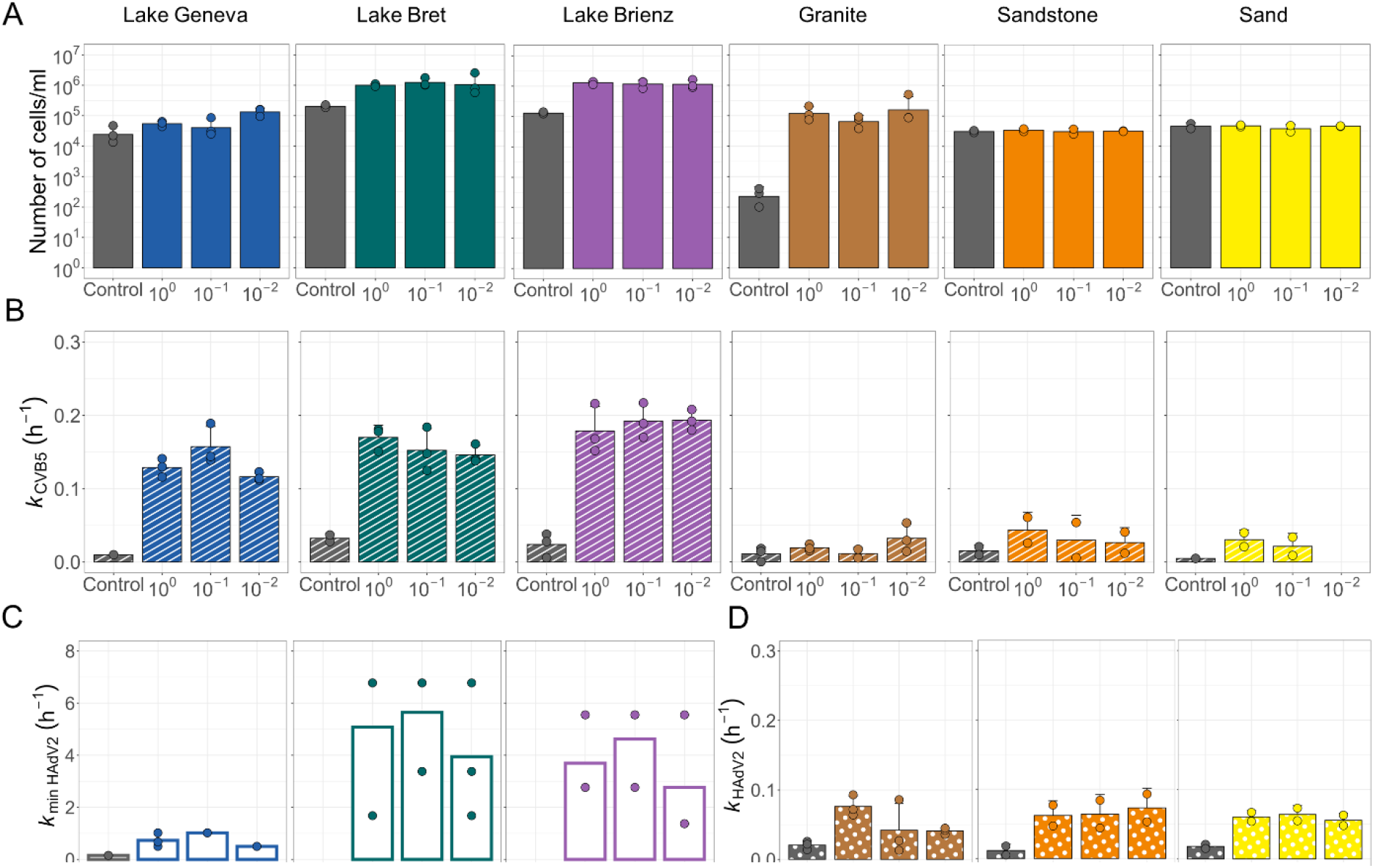
Cell density and viral inactivation in lakewater and groundwater chemostats. **(A)** Average cell densities of chemostat cultures within a day of viral spike (Day 5 for lakewater cultures; day 6 or 7 for groundwater cultures). Inactivation rate constants for CVB5 (kCVB5) **(B)** and HAdV2 (kHAdV2) **(C, D)** per water source. Because HAdV2 was rapidly inactivated in lakewater chemostats, minimal inactivation rate constants (kmin) were calculated for these chemostats using the time at which the virus concentration becomes undetectable (equation 2). This rates are indicated as empty bars. Error bars represent the standard deviation associated with k determined in replicate chemostats (n = 2-3).

For all chemostat cultures, we did not observe a significant correlation between seeding dilutions and calculated rate constants (**Figure S9**). Lastly, even though we observed bacterial growth in control chemostats, their viral inactivation was significantly lower compared to seeded chemostats (Welch’s two-sample t-test, *p* value < 0.05) for CVB5 in all lakewater chemostats and HAdV2 in groundwater chemostats (**Figure S10**). For CVB5 in groundwater, inactivation was comparable between controls and seeded chemostats as CVB5 showed greater stability in groundwater. Our chemostat cultures successfully grew and established freshwater communities with varying levels of antiviral activity against CVB5 and HAdV2. Despite the variation in diversity and community composition previously described, inactivation rate constants varied comparatively little among chemostats grown from the same source or freshwater type.

### Candidate biomarkers of virus inactivation in freshwater communities

Next, we aimed to identify candidate OTUs associated with viral inactivation. We used ANCOM-BC for differential abundance analyses. As microbial communities differed substantially among water types (**Figure 2A**), we conducted these analyses separately for groundwater and lakewater chemostats. For lakewater, we identified 107 taxa showing robust differential abundance in association with *k*_CVB5_ (q < 0.05), of which 99 were positively associated and 8 were negatively associated with inactivation (**Figure 4A**). As *k*_HAdV2_ could not be measured in lakewater, no biomarkers were identified for this virus. For groundwater communities, no taxa met the criteria for both statistical significance and robustness using either *k*_CVB5_ or *k*_HAdV2_. We therefore conducted an analysis using less stringent criteria to identify potentially meaningful associations that may require further validation. In this analysis, 797 OTUs showed significant associations with k_CVB5_ and 628 with *k*_HAdV2_ (q < 0.05). Of those, 373 OTUs were positively associated with k_CVB5_ and 340 with *k*_HAdV2_, (**Figure 4C and E**).

**Figure 4.**
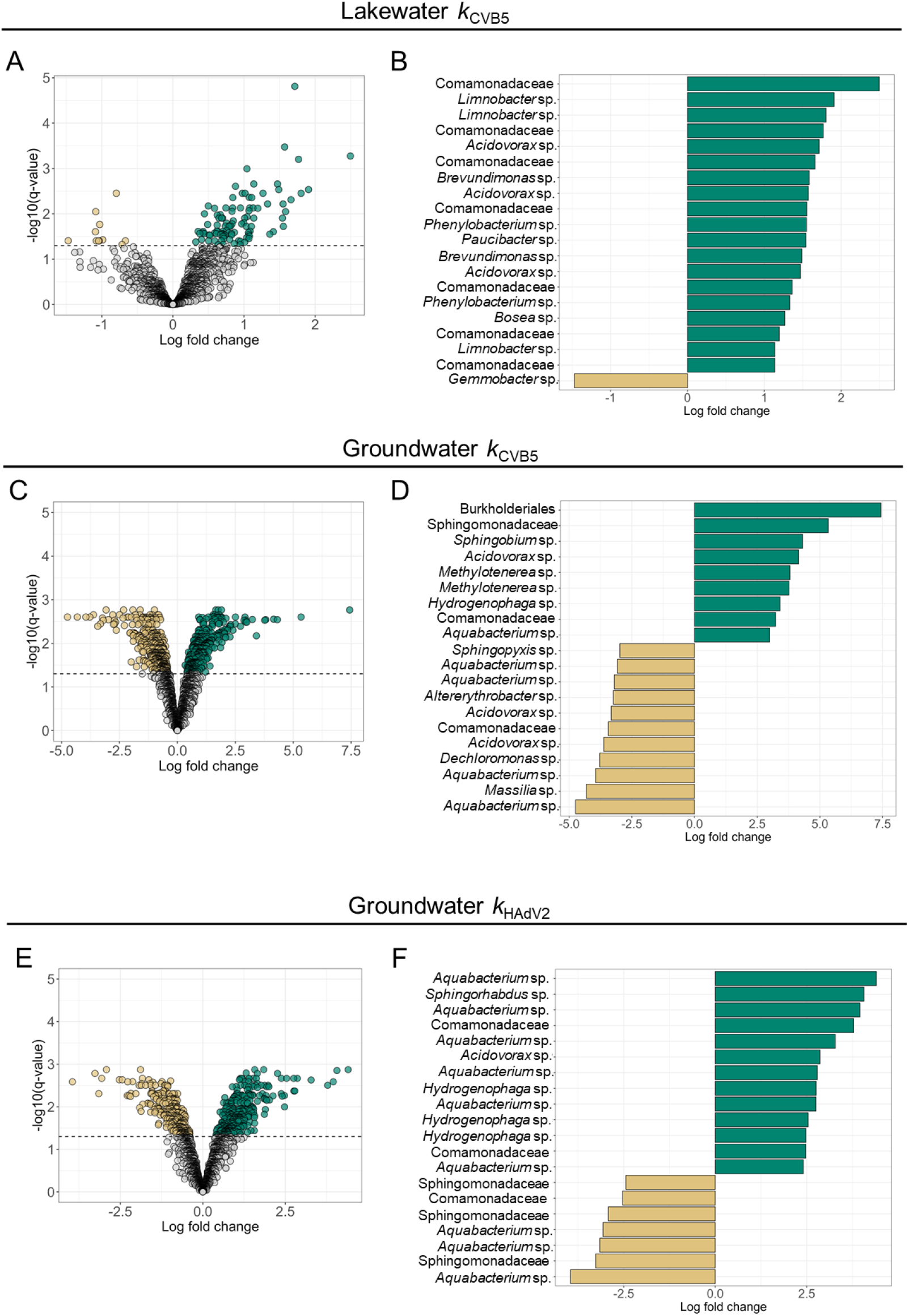
Differential abundance analyses for lakewater and groundwater chemostats. Analysis of Compositions of Microbiomes with Bias Correction (ANCOM-BC) was used to identify significant (q value < 0.05) bacteria operational taxonomic units (OTUs) associated with viral inactivation in chemostat cultures. Volcano plot of all differentially abundant taxa associated with k_CVB5_ found in lakewater chemostats **(A)** and in groundwater chemostats **(C)**, and with k_HAdV2_ found in groundwater chemostats **(E)**. Top 20 taxa with the largest estimated abundance changes as a function of k_CVB5_ in lakewater chemostats **(B)** and in groundwater chemostats **(D)**, and k_HAdV2_ in groundwater chemostats **(F)**.

Across both freshwater types, *Acidovorax* sp. were within the top 20 OTUs with the largest changes in relative abundance paralleled by changes in CVB5 inactivation (**Figure 4B, D and F**). This suggests faster inactivation of CVB5 in samples where *Acidovorax* sp. were relatively more abundant. In addition, in lakewater, abundances of *Limnobacter* sp., *Brevundimonas* sp. and *Phenylobacterium* sp. were positively associated with CVB5 inactivation, whereas in groundwater, *Methylotenerea* sp., *Aquabacterium* sp., and *Hydrogenophaga* sp. showed positive associations. In groundwater, we also observed several OTUs assigned to *Sphingopyxis* sp. that had positive associations with *k_CVB5_* and *k_HAdV2_*. Interestingly, particularly in groundwater, OTUs assigned to the same taxonomy sometimes showed opposing associations with viral inactivation. This may reflect strain-level differences in functional traits that are not resolved by taxonomic assignment. Differences in environmental context or other unmeasured factors may also contribute to these contrasting associations.

To assess whether the associations identified among the top 20 OTUs with the largest changes in abundance were consistent across source waters, we visually examined the relationship between *k* and OTU abundances. Some OTUs assigned to *Acidovorax* sp. were consistently abundant across lakewater chemostats and showed a positive association with *k*_CVB5_ (**Figure S11; Table S13**). In contrast, some associations appeared to be primarily driven by OTU abundances in Lake Brienz, including several OTUs assigned to *Brevundimonas* sp. and two *Phenylobacterium* sp. In groundwater, most associations were driven by OTU abundances in chemostats originating from the granite aquifer for the *k*_CVB5_ model, whereas associations in the *k*_HAdV2_ model were primarily driven by chemostats originating from the sand aquifer (**Figures S12–S13; Tables S14–S15**). This suggests that some taxa are consistently associated with viral inactivation across environments, whereas others show source- or virus-specific associations.

In summary, our analyses identified a set of candidate bacteria OTUs associated with viral inactivation. *Acidovorax* sp. was the main recurring taxon across both freshwater types. It was within the top 20 OTUs in lakewater and groundwater. In addition, in groundwater, OTUs assigned to *Hydrogenophaga* sp. and *Aquabacterium* sp. showed a similar association against calculated rate constants of both viruses. The consistent association of *Acidovorax* sp. with viral inactivation across different source waters and viral targets makes them promising candidates for further investigation. At the same time, source-specific candidate OTUs suggest that bacteria inactivation of viruses may be strongly influenced by the environmental context.

### Identifying predictors of viral inactivation in freshwater communities

After characterizing bacterial growth, diversity, and community composition and identifying candidate OTUs associated with viral inactivation, we next assessed if these biological parameters were predictive of viral inactivation. Multiple linear regression models were fitted separately for each freshwater type and virus. The models explained a substantial proportion of the variation in viral inactivation, with R² values ranging from 0.553 to 0.664, and all three models were statistically significant (*p* ≤ 0.022) (**Tables S16, S17 and S18**). However, the parameters associated with viral inactivation differed between freshwater types and viruses. In lakewater, the relative abundance of *Acidovorax* sp. was positively associated with *k*_CVB5_ (β = 0.0067, *p* value= 0.034), whereas the relative abundance of *Brevundimonas* sp. showed a positive but non-significant association (β = 0.02, *p* value = 0.073) (**Figure 5A**). In groundwater, *Sphingopyxis* sp. abundance was positively associated with both *k*_CVB5_ (β = 0.0200, *p* value= 0.0026) and *k*_HAdV2_ (β = 0.0085, *p* value= 0.023), while relative abundance of *Aquabacterium* sp. was positively associated only with *k*_CVB5_ (β = 0.0110, *p* value = 0.029) (**Figure 5B–C**).

**Figure 5.**
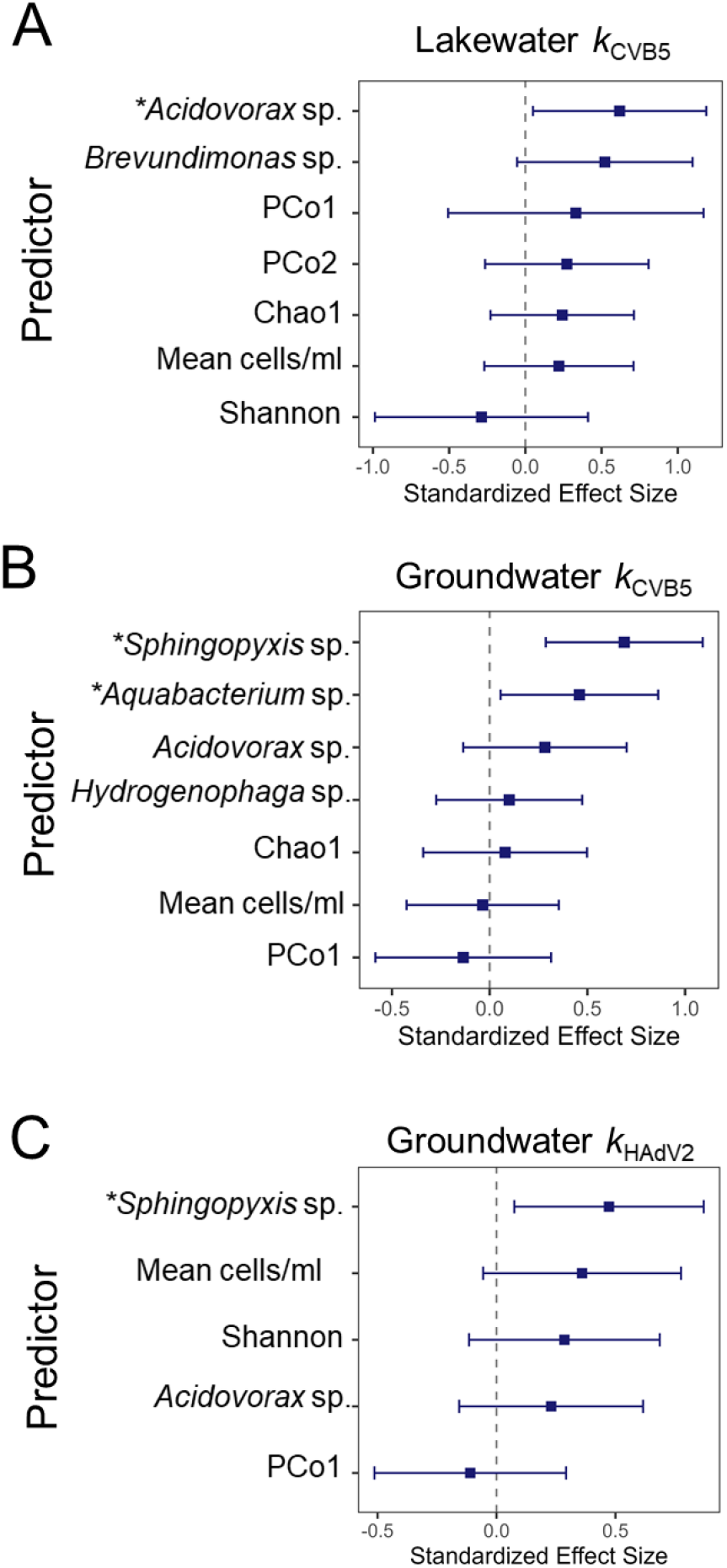
Predictors of viral inactivation in groundwater and lakewater chemostats. Multiple linear regression models were used to identify predictors of viral inactivation rate constants (k). Predictors included the relative abundance of selected bacterial taxa, bacterial cell counts, microbial community composition (PCo1 and PCo2), and bacterial diversity indices (Chao1 and Shannon). Values of k_CVB5_ were used as the response variable for lakewater **(A)** and groundwater communities **(B)**, and k_HAdV2_ for groundwater **(C)** communities. The dashed vertical line at zero indicates no association. Positive coefficients indicate that increasing values of the predictor are associated with increased inactivation, whereas negative coefficients indicate an inverse association.

In contrast, broader community-level parameters were not significant predictors of viral inactivation. Neither cell abundance, nor alpha diversity indices or community composition (PCo1 and PCo2) were significantly associated with inactivation in any of the models. Interestingly, in lakewater chemostats PCo1 showed positive associations with *k*_CVB5_ while in those from groundwater we observed negative associations for inactivation of both viruses. This indicates that groundwater communities which resemble lakewater communities had higher virus inactivation than groundwater communities that were more different to lakewater communities. All in all, these results suggest that variation in viral inactivation was more strongly associated with the abundance of specific bacterial OTUs than with overall bacterial abundance, alpha diversity, or broad differences in community composition.

## DISCUSSION

We implemented chemostat cultures to characterize viral inactivation in freshwater ecosystems. Our chemostat cultures successfully established freshwater communities with varying levels of antiviral activity. We observed 3 – 4 log_10_ reduction of CVB5 in lakewater and 1-2 log_10_ in groundwater after two days (**Figure S5 and S6**). For HAdV2, we observed > 4 log_10_ reduction after 8 hours in lakewater (**Figure S7**) and 2 log_10_ after two days in groundwater (**Figure S8**). Previous research using batch cultures reported lower inactivation compared to our systems. Romanenko *et al*. (2025) and Meibom *et al.* (2026), reported between 1 - 2 log_10_ infectivity decrease of CVB5 after 48 hours in microcosms from Lake Geneva ^13,22^. Another study assessing enteric virus persistence in batch reactors of river water reported that the average time for 1 log_10_ unit infectivity loss of two enteric viruses in microcosms was 13 days for HAdV41 and 11 days for CVB2, and both viruses were still detected after 70 days ^23^. In addition, Ogorzaly *et al.* (2010), tested the inactivation of HAdV2 in batch groundwater microcosms and found 1 log_10_ unit infectivity decrease after 20 days and virus was still detected after 120 days ^24^. Although these studies used different viruses and source waters from those investigated here, our data suggest that chemostat cultures promoted viral inactivation in lakewater and groundwater. This may reflect differences in bacterial physiological state, as chemostats maintain bacteria under steady-state growth conditions, unlike batch cultures. It may also reflect fundamental differences in traits of bacteria selected under chemostat conditions. For instance, the constant but low-nutrient conditions simulated by our chemostat systems is expected to foster slow-but-efficient bacterial metabolic strategies over fast-but-inefficient growth^25,26^. Hence, bacterial traits linked to efficient resource use may be associated with virus inactivation, particularly in diluted habitats such as groundwater or lake ecosystems. Together, these findings suggest that both bacterial physiology and community composition may influence viral inactivation.

We aimed to generate freshwater microbial communities with gradients in diversity and composition to better understand the influence of bacterial diversity on viral inactivation. Dilution-to-extinction is thought to remove rare species and decrease bacterial richness^27^. Here, dilution-to-extinction exerted a modest effect on Chao1 and Shannon indices in lakewater chemostats, but a significant decrease in chemostats from granite and sandstone aquifers (**Figure 2B and 2C**). This is consistent with previous observations of greater diversity gradients in groundwater compared to surface water cultures^14^. Bray-Curtis dissimilarity showed that community composition shifted across all starting dilutions in both lakewater and groundwater chemostats (**Figure 2D**), although these shifts were smaller than the differences between the two water types. Together, these results indicate that dilution-to-extinction altered not only richness and evenness of the communities but also the identity of the bacteria established in the chemostats.

Although dilution-to-extinction produced different communities, viral inactivation varied comparatively little among most chemostats grown from the same source or freshwater type. Notably, all chemostats showed significantly greater viral inactivation than our controls, indicating that the observed inactivation was driven by microbial activity rather than differences in physical or chemical parameters among treatments. We did not observe a strong association between Chao1 and Shannon indices and viral inactivation in lakewater. In contrast, Romanenko *et al.* (2025), reported a positive association between microbial diversity in lakewater and antiviral activity against echovirus 11^13^. In the present study, we observed a similar effect only for groundwater chemostat communities from sandstone and sand aquifers; these chemostats showed a modest reduction in CVB5 inactivation in cultures with greater richness (**Figure 3B**). This is likely because we generated greater gradients down to lower richness values in groundwater compared to lakewater chemostats. Other studies have successfully reduced richness of freshwater bacterial communities and observed contrasting effects on community function. Mao *et al.* (2023) found that dilution of lakewater removed copiotrophs and rare taxa and these changes altered carbon utilization ^28^, whereas Peter *et al.* (2011) found that although dilution reduced richness, rates of chitin and cellulose degradation were mostly explained by community composition rather than by richness itself ^29^. Together, these findings suggest that changes in freshwater community composition, rather than richness or evenness alone, are more important in determining bacterial-mediated viral inactivation.

We found *Acidovorax* sp. enriched in some of our freshwater chemostat cultures and differential abundance analyses identified abundance of some OTUs from this genus as correlated with viral inactivation (**Figure 2D and Figure 4**). The genus *Acidovorax* comprises gram-negative bacteria that are widely distributed in diverse environments including freshwater ecosystems^30,31^. Comparative genomic analyses by Siani *et al.* (2021), showed that both plant-beneficial and pathogenic *Acidovorax* sp. strains produce hydrolytic enzymes and a range of secondary metabolites, including antimicrobial compounds, organic acids, and phytohormones^32^. In addition, several studies have reported that *Acidovorax* species produce extracellular polymeric substances, form biofilms and degrade polycyclic aromatic compounds ^33–35^. We also identified OTUs from the genus *Sphingopyxis* to be positively associated with viral inactivation in groundwater and the abundance of these OTUs was a significant predictor of the inactivation of CVB5 and HAdV2. This genus comprises bacteria known for the biodegradation capabilities against environmental contaminants^36^. They also secrete peptidases and proteases, including serine proteases ^37^, whose production we have previously seen associated with viral inactivation^11^. Exopolysaccharide and biofilm production, and protease secretion have been previously described in other organisms as mechanisms of viral inactivation^8,38,39^. All in all, these traits could facilitate virus adsorption or retention within microbial aggregates, thereby prolonging exposure to bacteria, and providing a potential mechanism link of *Acidovorax* sp. and *Sphingopyxis* sp. to viral inactivation.

Other identified candidate biomarkers of viral inactivation were OTUs from the genera *Aquabacterium* sp. and *Phenylobacterium* sp. (**Figure 4**). *Aquabacterium* sp. have the metabolic potential to degrade complex hydrocarbon substrates and other organic contaminants ^40,41^, however, little is known about the potential roles of these genera in virus–microbe interactions. In earlier work, we identified an association between *Chitinophaga* sp. abundance and inactivation of echovirus 11 in lakewater ^13^. Here, abundance analyses of chemostat cultures did not identify OTUs from this genus to be associated with either *k*_CVB5_ or *k*_HAdV2_. This discrepancy reflects the selective pressures specific to our chemostat culture conditions ^14^ or differences in inactivation mechanisms against echovirus 11 compared to the viruses used here. Together, our findings identified bacterial taxa that are consistently associated with viral inactivation of CVB5 and HAdV2 and represent promising candidate biomarkers. Further research using representative isolates or enrichment cultures for these genera are needed to determine the mechanisms underlying viral inactivation and to establish whether these taxa play a direct role in the process.

Chemostats provide multiple advantages for the study of freshwater communities compared to batch cultures (e.g., constant replenishment of resources and the washout of waste products). However, they still have important limitations. Some of the chosen experimental conditions in this study (e.g., flowrates, aeration, incubation in the dark) reduced freshwater communities’ complexity. These conditions were previously optimized to manipulate diversity and support the growth of lakewater and groundwater communities ^14^. Our goal was not to fully replicate natural freshwater conditions but rather to establish communities with diversity gradients from different source freshwaters. Consequently, the candidate bacterial biomarkers for viral inactivation here identified should be further validated in natural lakewater and groundwater systems. Assessing their abundance in these environments and their association with viral inactivation will determine their ecological relevance. Another limitation of our experimental design was that we were unable to avoid growth in our chemostat controls. Previous studies culturing freshwater communities have used multiple rounds of autoclaving to eliminate bacteria spores ^29^. However, we chose not to autoclave our freshwater as this process can alter or degrade some of the nutrients present and change important chemical parameters including pH. Interestingly, despite the unintended growth in our chemostat controls, we found that the bacteria grown in our controls did not inactivate the viruses spiked into the system. This observation further supports our hypothesis that viral inactivation is associated with the presence of key taxa. Lastly, the range of bacterial richness achieved in this study was narrower than that in our previous work, in which viral inactivation showed a saturating relationship versus richness^13^. The relatively high richness of the communities established here may therefore have limited our ability to detect an effect of richness on viral inactivation, as the communities may have already been within this saturation range.

Overall, by comparing viral inactivation and bacterial community composition in chemostat cultures derived from multiple lake and groundwaters, we provide new insights into the role of freshwater bacterial communities in shaping the environmental stability of human enteric viruses. The identification of bacterial taxa from diverse genera that were consistently associated with viral inactivation suggests that this capacity is mediated by specialized microorganisms. Our work provides a deeper understanding of the microbial processes that naturally limit viral stability in freshwater.

## Acknowledgements

This work was supported by the Swiss National Science Foundation (grant no. 215226). We thank Jessica Sherman (USDA-ARS) and the Central Environmental Laboratory (GR-CEL-EPFL) for evaluating water chemistry in lakewater and groundwater samples. Thanks also to Sarah Opelt (USDA-ARS) for assistance with virus propagation and enumeration. Any use of trade, firm, or product names is for descriptive purposes only and does not imply endorsement by the U.S. Government. USDA is an equal opportunity provider and employer.

## Supporting Information

Additional supporting information: methods, bacterial growth and inactivation curves, calculated diversity indices, microbial abundances analyses results and multiple linear regression models details.

## Data availability

All data (virus titres and bacteria abundances in chemostats cultures) are available on Zenodo under the following link: 10.5281/zenodo.23495094. Sequencing data is available in National Center for Biotechnology Information (NCBI) – Sequence Read Archive (SRA) under the following accession number: PRJNA1549321.

