## Supplemental Figures and Tables for "Freshwater Microbiomes Shape Viral Inactivation in Continuous Cultures"

Table S12. Inactivation rate constants for groundwater chemostats against Human Adenovirus 2 (HAdV2) .7

Table S16. Multiple linear regression coefficients of $k_{CVB5}$ against biological parameters for LW chemostats 7

Table S17. Multiple linear regression coefficients of $k_{CVB5}$ against biological parameters for GW chemostats8

**Supplemental Method S1. Sequencing data processing and microbial diversity analyses.** The sequencing .pod5 files were basecalled post hoc using Dorado (v0.9.1, super high accuracy) and FASTQ files from all experiments were used as combined input into the MetONTIIME workflow v2.1.0 [1] with Nextflow v 24.10.5 [2]. The MetONTIIME workflow parameters were as follows: Maximum number of reads per sample = 75000, Read Length = 1400 – 1700, Minimum Quality of Reads: 18, Clustering identity: 0.95, Minimum Consensus = 0.7, Minimum Query Coverage = 0.8, and Minimum Identity of 0.9. Sequences were taxonomically assigned against SILVA 16S rRNA database (version 138.2). Taxonomy and OTU table outputs were then analyzed in R. Bacterial OTUs detected in the extraction blanks were identified as potential contaminants based on quantitative comparisons of their abundances between extraction blanks and experimental samples using the *decon* function from the R package *microDecon* (version 1.0.2)[3], and were subsequently removed from the species abundance table. After decontamination, bacterial OTUs that were found once or had only had one read across samples in each experiment were removed. Microbial community analyses were performed using the *microeco* R package (version 2.0.0)[4].

**Table S1. Physicochemical parameters measured on sampling day from each water source**

| Parameters | Lake Geneva | Lake Bret | Lake Brienz | Granite | Sandstone | Sand |
|---|---|---|---|---|---|---|
| Date | Feb (2025) | Oct (2025) | Dec (2025) | May (2025) | Dec (2025) | Dec (2025) |
| Temperature (°C) | 8.2 | 15.4 | 8.4 | - | - | - |
| pH | 7.8 | 8.13 | 8.04 | 6.46 | 7.87 | 7.11 |
| Dissolved Oxygen (mg/L) | 10.7 | 9.03 | 11.01 | - | - | - |
| Conductivity (μS/cm) | 319 | 293 | 173.6 | 292 | 411 | 382 |
| [1]TOC (mg/L) | 0.500 | 1.664 | < 0.25 | 42.2 | 0.633 | 0.637 |
| [2]TN (mg/L) | 0.631 | < 0.5 | 0.290 | 5.83 | 2.44 | 0.0075 |

*[1]Total Organic Carbon; [2]Total Nitrogen*

**Table S2. Chao1 and Shannon indexes calculated from lakewater and groundwater communities before chemostat cultures**

| | Chao1 | Shannon |
|---|---|---|
| Lake Geneva | 7023 | 6.07 |
| Lake Bret | 5889 | 6.38 |
| Lake Brienz | 4436 | 6.31 |
| Sand | 2816 | 5.35 |
| Sandstone | 3886 | 5.80 |
| Granite | 4879 | 7.08 |

**Table S3. Pairwise Bray-Curtis dissimilarities between different source waters**

| Water Type | Water Source 1 | Water Source 2 | Mean Bray-Curtis | SD Bray-Curtis |
|---|---|---|---|---|
| Groundwater | Granite | Sandstone | 0.89 | 0.09 |
| | Granite | Sand | 0.95 | 0.03 |
| | Sand | Sandstone | 0.97 | 0.01 |
| Lakewater | Geneva | Bret | 0.85 | 0.10 |
| | Geneva | Brienz | 0.83 | 0.15 |
| | Brienz | Bret | 0.90 | 0.10 |

**Table S4. Pairwise Bray-Curtis dissimilarities between different starting dilutions**

| Water Type | Water Source | Condition 1 | Condition 2 | Mean Bray-Curtis | SD Bray-Curtis |
|---|---|---|---|---|---|
| Groundwater | Granite | $10^{0}$ | $10^{-1}$ | 0.69 | 0.07 |
| | | $10^{0}$ | $10^{-2}$ | 0.77 | 0.08 |
| | | $10^{-2}$ | $10^{-1}$ | 0.66 | 0.17 |
| | Sandstone | $10^{0}$ | $10^{-1}$ | 0.64 | 0.12 |
| | | $10^{0}$ | $10^{-2}$ | 0.83 | 0.09 |
| | | $10^{-2}$ | $10^{-1}$ | 0.81 | 0.12 |
| | Sand | $10^{0}$ | $10^{-1}$ | 0.79 | 0.02 |
| | | $10^{0}$ | $10^{-2}$ | 0.82 | 0.05 |
| | | $10^{-2}$ | $10^{-1}$ | 0.81 | 0.04 |
| Lakewater | Geneva | $10^{0}$ | $10^{-1}$ | 0.70 | 0.08 |

| | | | | | |
|---|---|---|---|---|---|
| | | $10^0$ | $10^{-2}$ | 0.71 | 0.08 |
| | | $10^{-2}$ | $10^{-1}$ | 0.60 | 0.08 |
| | Bret | $10^0$ | $10^{-1}$ | 0.77 | 0.12 |
| | | $10^0$ | $10^{-2}$ | 0.87 | 0.10 |
| | | $10^{-2}$ | $10^{-1}$ | 0.81 | 0.13 |
| | Brienz | $10^0$ | $10^{-1}$ | 0.89 | 0.04 |
| | | $10^0$ | $10^{-2}$ | 0.92 | 0.03 |
| | | $10^{-2}$ | $10^{-1}$ | 0.72 | 0.03 |

**Table S5. Chao1 coefficient for lakewater chemostats**

| Condition | Lake Geneva | Lake Bret | Lake Brienz |
|---|---|---|---|
| $10^0$ | 3808 | 4457 | 3007 |
| | 3873 | 6039 | 3030 |
| | 5205 | 5448 | 4009 |
| Mean ± SD | 4295 ± 754 | 5314 ± 804 | 3348 ± 291 |
| $10^{-1}$ | 2484 | 3033 | 3607 |
| | 3533 | 3279 | 4012 |
| | 3561 | 3433 | 4184 |
| Mean ± SD | 3192 ± 601 | 3248 ± 203 | 3934 ± 291 |
| $10^{-2}$ | 3094 | 3856 | 2901 |
| | 3634 | 3647 | 2596 |
| | 2692 | 3449 | 2808 |
| Mean ± SD | 3140 ± 471 | 3650 ± 219 | 2768 ± 159 |

**Table S6. Chao1 coefficient for groundwater chemostats**

| Condition | Granite | Sandstone | Sand |
|---|---|---|---|
| $10^0$ | 3966 | 4068 | 2620 |
| | 3628 | 4098 | 3192 |
| | 2806 | - | - |
| Mean ± SD | 3467 ± 597 | 4083 ± 21 | 2906 ± 404 |
| $10^{-1}$ | 2468 | 2702 | 1920 |
| | 2747 | 3265 | 2293 |
| | 2126 | - | - |
| Mean ± SD | 2447 ± 311 | 2984 ± 398 | 2107 ± 264 |
| $10^{-2}$ | 654 | 2772 | 1842 |
| | 1543 | 2449 | 2648 |
| | 1712 | - | - |
| Mean ± SD | 1303 ± 568 | 2611 ± 228 | 2245 ± 570 |

**Table S7. Shannon coefficient for lakewater chemostats**

| Condition | Lake Geneva | Lake Bret | Lake Brienz |
|---|---|---|---|
| $10^0$ | 4.76 | 4.88 | 2.64 |
| | 4.02 | 5.4 | 2.79 |
| | 4.65 | 5.51 | 3.52 |
| Mean ± SD | 4.48 ± 0.39 | 5.26 ± 0.34 | 2.98 ± 0.46 |

| | | | |
|---|---|---|---|
| $10^{-1}$ | 3.35 | 4.21 | 5.09 |
| | 3.7 | 3.71 | 4.99 |
| | 3.9 | 4.3 | 4.87 |
| Mean ± SD | 3.65 ± 0.28 | 4.07 ± 0.31 | 4.98 ± 0.11 |
| $10^{-2}$ | 3.51 | 4.88 | 4.85 |
| | 4.33 | 5.08 | 4.31 |
| | 3.88 | 4.65 | 4.48 |
| Mean ± SD | 3.91 ± 0.41 | 4.87 ± 0.10 | 4.55 ± 0.28 |

**Table S8. Shannon coefficient for groundwater chemostats**

| Condition | Granite | Sandstone | Sand |
|---|---|---|---|
| $10^{0}$ | 5.03 | 4.75 | 4.89 |
| | 5.14 | 5.31 | 5.13 |
| | 4.04 | - | - |
| Mean ± SD | 4.74 ± 0.60 | 5.03 ± 0.40 | 5.01 ± 0.17 |
| $10^{-1}$ | 3.76 | 2.75 | 4.39 |
| | 4.33 | 3.97 | 3.58 |
| | 3.86 | - | - |
| Mean ± SD | 3.98 ± 0.31 | 3.36 ± 0.86 | 3.99 ± 0.57 |
| $10^{-2}$ | 1.82 | 4.14 | 3.92 |
| | 2.99 | 3.6 | 3.74 |
| | 3.4 | - | - |
| Mean ± SD | 2.74 ± 0.81 | 3.87 ± 0.38 | 3.83 ± 0.13 |

**Table S9. Inactivation rate constants for lakewater chemostats against Coxsackievirus B5 (CVB5)**

| | Lake Geneva | | Lake Bret | | Lake Brienz | |
|---|---|---|---|---|---|---|
| **Condition** | ***k*** **($h^{-1}$)** | **95% CI** | ***k*** **($h^{-1}$)** | **95% CI** | ***k*** **($h^{-1}$)** | **95% CI** |
| $10^{0}$ | 0.12 | (0.08 – 0.15) | 0.18 | (0.12 – 0.24) | 0.15 | (0.09 – 0.22) |
| | 0.14 | (0.08 – 0.20) | 0.15 | (0.09 – 0.21) | 0.17 | (0.08 – 0.26) |
| | 0.13 | (0.10 – 0.16) | 0.18 | (0.11 – 0.25) | 0.22 | (0.14 – 0.29) |
| $10^{-1}$ | 0.14 | (0.10 – 0.18) | 0.18 | (0.15 – 0.22) | 0.17 | (0.09 – 0.25) |
| | 0.14 | (0.11 – 0.18) | 0.12 | (0.08 – 0.17) | 0.19 | (0.12 – 0.26) |
| | 0.19 | (0.15 – 0.23) | 0.15 | (0.10 – 0.20) | 0.22 | (0.15 – 0.29) |
| $10^{-2}$ | 0.12 | (0.06 – 0.19) | 0.16 | (0.09 – 0.23) | 0.19 | (0.12 – 0.7) |
| | 0.11 | (0.06 – 0.16) | 0.14 | (0.06 – 0.22) | 0.18 | (0.1 – 0.26) |
| | 0.11 | (0.09 – 0.14) | 0.14 | (0.06 – 0.21) | 0.21 | (0.14 – 0.28) |

**Table S10. Inactivation rate constants for groundwater chemostats against Coxsackievirus B5 (CVB5)**

| | Granite | | Sandstone | | Sand | |
|---|---|---|---|---|---|---|
| **Condition** | ***k*** **($h^{-1}$)** | **95% CI** | ***k*** **($h^{-1}$)** | **95% CI** | ***k*** **($h^{-1}$)** | **95% CI** |
| $10^{0}$ | 0.01 | (-0.01 – 0.03) | 0.06 | (0.05 – 0.07) | 0.04 | (0.02 – 0.06) |
| | 0.02 | (-0.01 – 0.06) | 0.03 | (0.02 – 0.03) | 0.02 | (0 – 0.03) |
| | 0.02 | (0 – 0.04) | - | - | - | - |
| $10^{-1}$ | 0.02 | (-0.01 – 0.05) | 0.01 | (0 – 0.02) | 0.01 | (-0.01 – 0.02) |
| | 0.01 | (-0.02 – 0.03) | 0.05 | (0.03 – 0.07) | 0.03 | (0 – 0.05) |
| | 0 | (-0.02 – 0.02) | - | - | - | - |

| | | | | | | |
|---|---|---|---|---|---|---|
| $10^{-2}$ | 0.05 | (0.03 – 0.08) | 0.01 | (0 – 0.02) | -0.01 | (-0.03 – 0.01) |
| | 0.01 | (0 – 0.03) | 0.04 | (0.03 – 0.05) | 0 | (-0.02 – 0.02) |
| | 0.03 | (-0.01 – 0.06) | - | - | - | - |

**Table S11. Minimal inactivation rate constants ($k_{min}$; Equation 2) for lakewater chemostats against Human Adenovirus 2 (HAdV2)**

| | Lake Geneva | Lake Bret | Lake Brienz |
|---|---|---|---|
| **Condition** | $k_{min}$ ($h^{-1}$) | | |
| $10^0$ | 0.54 | 7.12 | 5.90 |
| | 0.73 | 7.12 | 2.94 |
| | 1.11 | 1.77 | 2.94 |
| $10^{-1}$ | 1.11 | 3.55 | 2.94 |
| | 1.11 | 7.12 | 5.90 |
| | 1.11 | 7.12 | 5.90 |
| $10^{-2}$ | 0.54 | 7.12 | 5.90 |
| | 0.54 | 3.55 | 1.46 |
| | 0.54 | 1.77 | 1.46 |

**Table S12. Inactivation rate constants for groundwater chemostats against Human Adenovirus 2 (HAdV2)**

| | Granite | | Sandstone | | Sand | |
|---|---|---|---|---|---|---|
| **Condition** | $k$ ($h^{-1}$) | 95% CI | $k$ ($h^{-1}$) | 95% CI | $k$ ($h^{-1}$) | 95% CI |
| $10^0$ | 0.09 | (0.08 – 0.11) | 0.08 | (0.06 – 0.09) | 0.07 | (0.06 – 0.08) |
| | 0.06 | (0.04 – 0.09) | 0.05 | (0.04 – 0.06) | 0.05 | (0.03 – 0.07) |
| | 0.07 | (0.05 – 0.09) | - | - | - | - |
| $10^{-1}$ | 0.03 | (0.01 – 0.05) | 0.04 | (0.03 – 0.06) | 0.07 | (0.06 – 0.09) |
| | 0.09 | (0.06 – 0.11) | 0.08 | (0.07 – 0.1) | 0.06 | (0.03 – 0.08) |
| | 0.01 | (-0.02 – 0.04) | - | - | - | - |
| $10^{-2}$ | 0.04 | (0 – 0.07) | 0.05 | (0.04 – 0.07) | 0.06 | (0.04 – 0.09) |
| | 0.04 | (0.02 – 0.06) | 0.09 | (0.08 – 0.11) | 0.05 | (0.03 – 0.07) |
| | 0.05 | (0.02 – 0.07) | - | - | - | - |

**Table S16. Multiple linear regression coefficients of $k_{CVB5}$ against biological parameters for LW chemostats**

| | Beta coefficients (β) | Std. Error | t value | P value |
|---|---|---|---|---|
| Chao1 | $8.82 \times 10^{-6}$ | $8.2 \times 10^{-6}$ | 1.02 | 0.32 |
| Mean cells | 0.0105 | 0.0112 | 0.941 | 0.359 |
| Shannon | -0.0121 | 0.140 | -0.862 | 0.399 |
| PCo1 | 0.0363 | 0.0439 | 0.827 | 0.418 |
| PCo2 | 0.0363 | 0.0361 | 1.06 | 0.302 |
| *Acidovorax* relative abundances | 0.0067 | 0.003 | 2.28 | 0.0345 |
| *Brevundimonas* relative abundances | 0.0224 | 0.012 | 1.9 | 0.073 |

Residual standard error: 0.0219 on 19 degrees of freedom
Multiple R-squared: 0.642, Adjusted R-squared: 0.510
F-statistic: 4.87 on 7 and 19 DF, $p$ value: 0.002766

**Table S17. Multiple linear regression coefficients of $k_{CVB5}$ against biological parameters for GW chemostats**

| | **Beta coefficients (β)** | **Std. Error** | **t value** | **P value** |
|---|---|---|---|---|
| Chao1 | $1.7 \times 10^{-6}$ | $4.2 \times 10^{-6}$ | 0.41 | 0.69 |
| Mean cells | −0.0033 | 0.017 | −0.20 | 0.85 |
| PCo1 | −0.0088 | 0.014 | −0.65 | 0.53 |
| *Acidovorax* relative abundances | 0.0062 | 0.0043 | 1.46 | 0.17 |
| *Sphingopyxis* relative abundances | 0.02 | 0.0053 | 3.71 | 0.0026 |
| *Aquabacterium* relative abundances | 0.011 | 0.0044 | 2.46 | 0.029 |
| *Hydrogenophaga* relative abundances | 0.0023 | 0.0039 | 0.58 | 0.57 |

Residual standard error: 0.01344 on 13 degrees of freedom
Multiple R-squared: 0.6643, Adjusted R-squared: 0.4836
F-statistic: 3.675 on 7 and 13 DF, *p* value: 0.0206

**Table S18. Multiple linear regression coefficients of $k_{HAdV2}$ against biological parameters for GW chemostats**

| | **Beta coefficients (β)** | **Std. Error** | **t value** | **P value** |
|---|---|---|---|---|
| Shannon | 0.0070 | 0.0046 | 1.5 | 0.15 |
| Mean cells | 0.038 | 0.021 | 1.8 | 0.085 |
| PCo1 | −0.0083 | 0.014 | −0.58 | 0.57 |
| *Acidovorax* relative abundances | 0.00051 | 0.00040 | 1.3 | 0.22 |
| *Sphingopyxis* relative abundances | 0.0085 | 0.0034 | 2.5 | 0.023 |

Residual standard error: 0.01654 on 15 degrees of freedom
Multiple R-squared: 0.5529, Adjusted R-squared: 0.4039
F-statistic: 3.71 on 5 and 15 DF, *p* value: 0.02193

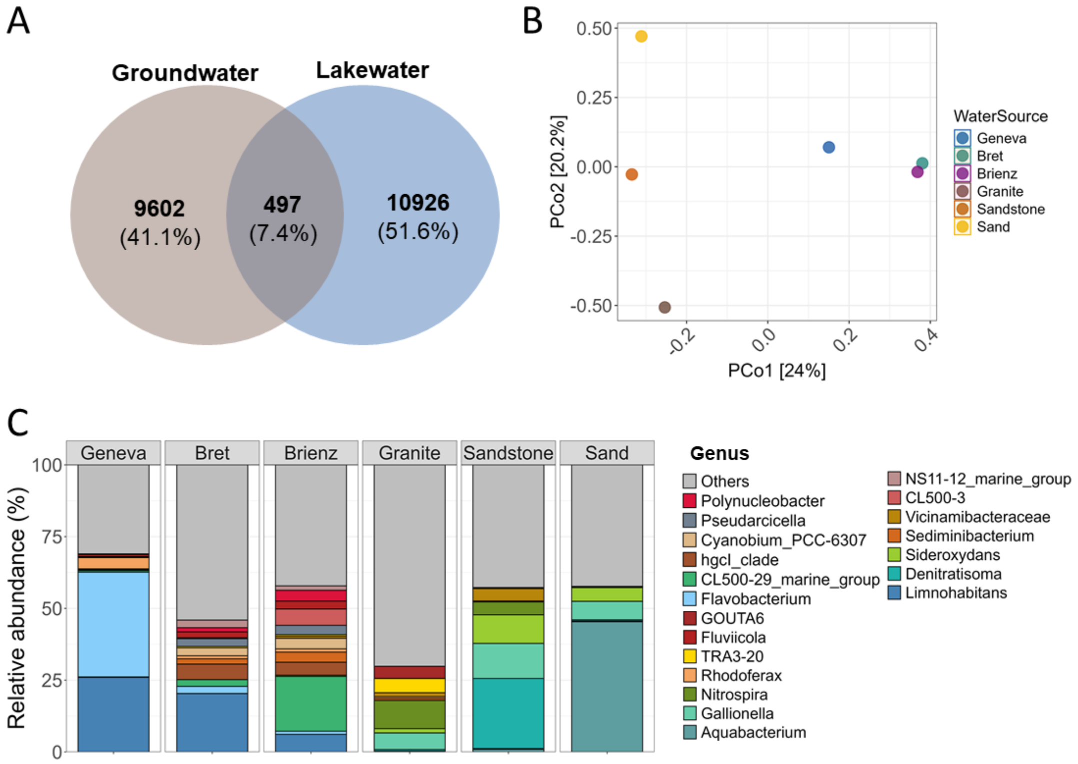

***Figure S1. Bacterial communities across water sources before chemostat cultures. (A)*** *Venn diagram with shared and unique OTUs between lake water and groundwater samples.* ***(B)*** *Principal coordinate analysis (PCoA) of Bray–Curtis distances calculated from communities.* ***(C)*** *Relative abundance of the top 20 most abundant bacteria genera found on each source.*

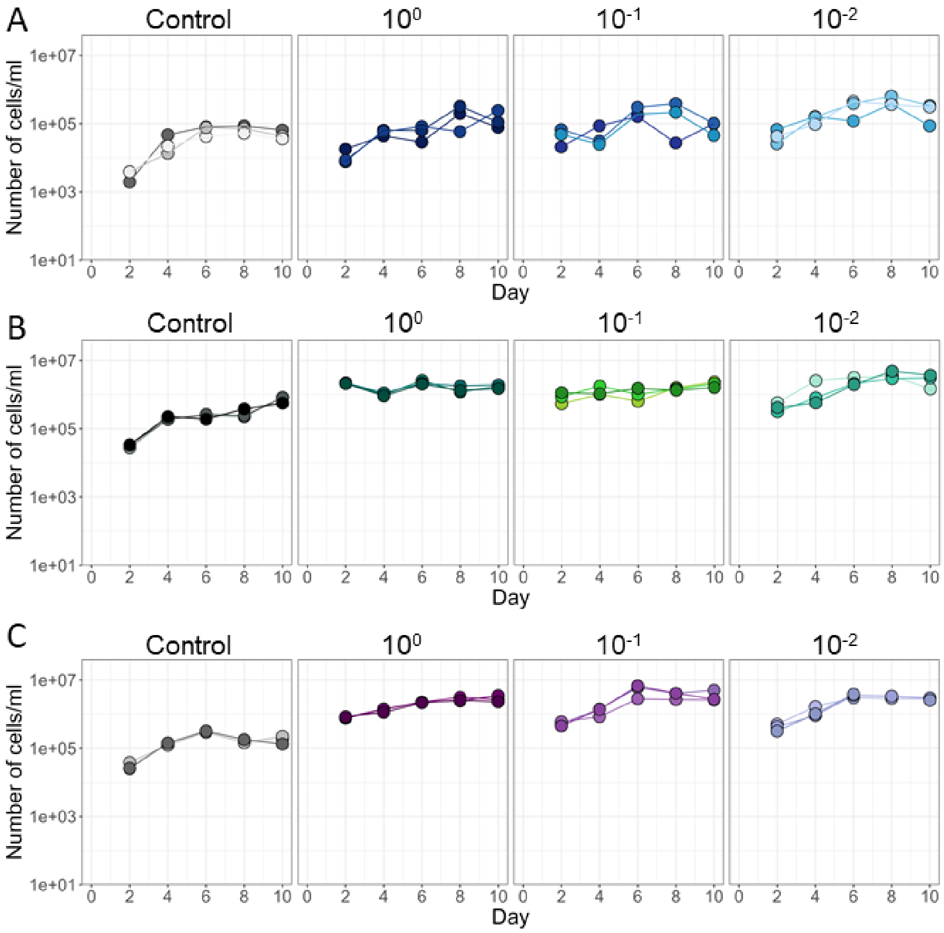

***Figure S2. Growth inside lakewater chemostats.*** *Undiluted ($10^0$) or diluted ($10^{-1}$ or $10^{-2}$) water from Lake Geneva **(A)**, Lake Bret **(B)** and Lake Brienz **(C)** was seeded into chemostat cultures and regrown for 10 days. Samples were taken every other day from the effluent and fixed with 3.7 % formaldehyde. To estimate the number of cells/ml per culture, samples were stained with SYTO13 and analysed by flow cytometry. As a control, sterile (filtered and UV-treated) water from each source was seeded and spiked with 1% penicillin and streptomycin every other day. Three biological replicates were grown per condition (n=3).*

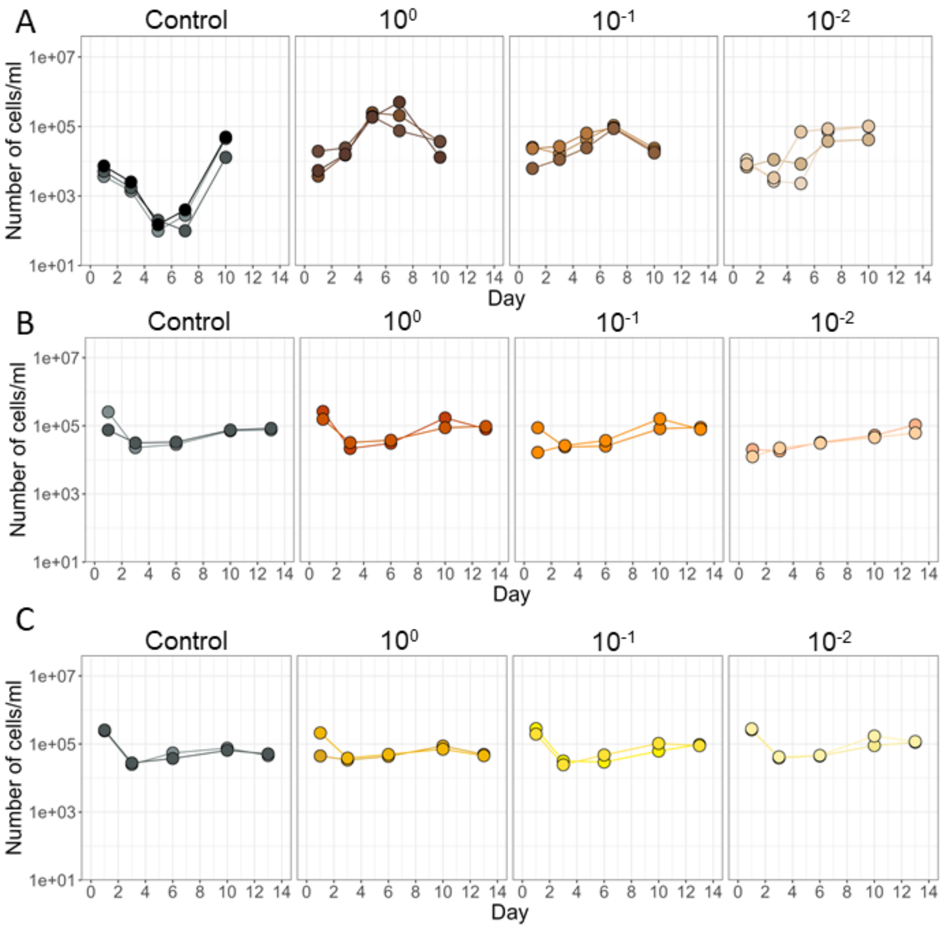

***Figure S3. Growth inside groundwater chemostats.*** *Undiluted ($10^0$) or diluted ($10^{-1}$ or $10^{-2}$) from granite* ***(A)****, sandstone* ***(B)*** *and sand* ***(C)*** *aquifers was seeded into chemostat cultures and regrown for 10-14 days. Samples were taken every 2-3 days from the effluent and fixed with 3.7 % formaldehyde. To estimate the number of cells/ml per culture, samples were stained with SYTO16 and analysed by flow cytometry. As a control, sterile (filtered and UV-treated) water from each source was seeded and spiked with 1% penicillin and streptomycin. Two or three biological replicates were grown per condition (n = 2-3).*

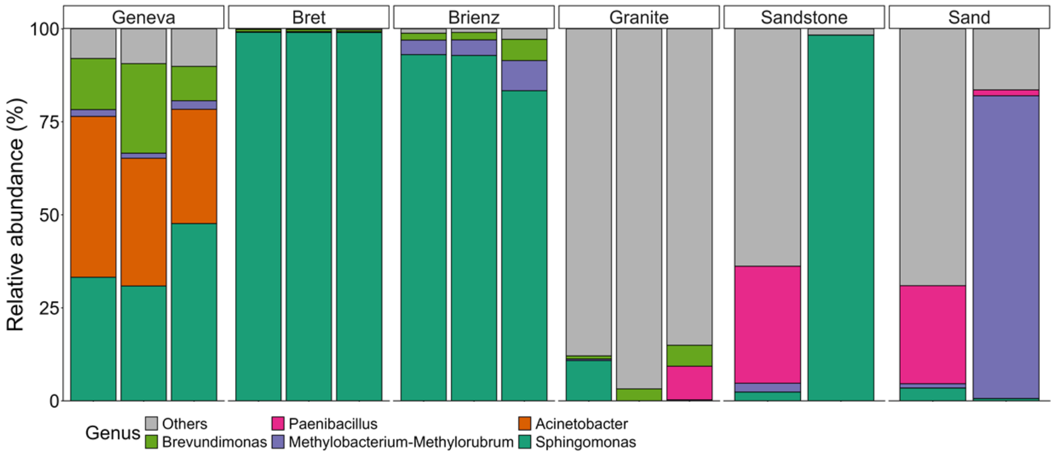

***Figure S4. Relative abundance of bacteria genera on unseeded controls.*** *Two to three unseeded controls were maintained alongside seeded chemostats throughout experiments. Despite the use of antibiotics every other day to avoid bacterial growth, we observed growth in these chemostats. However, inactivation profiles and community composition were different in these chemostats compared to experimental ones.*

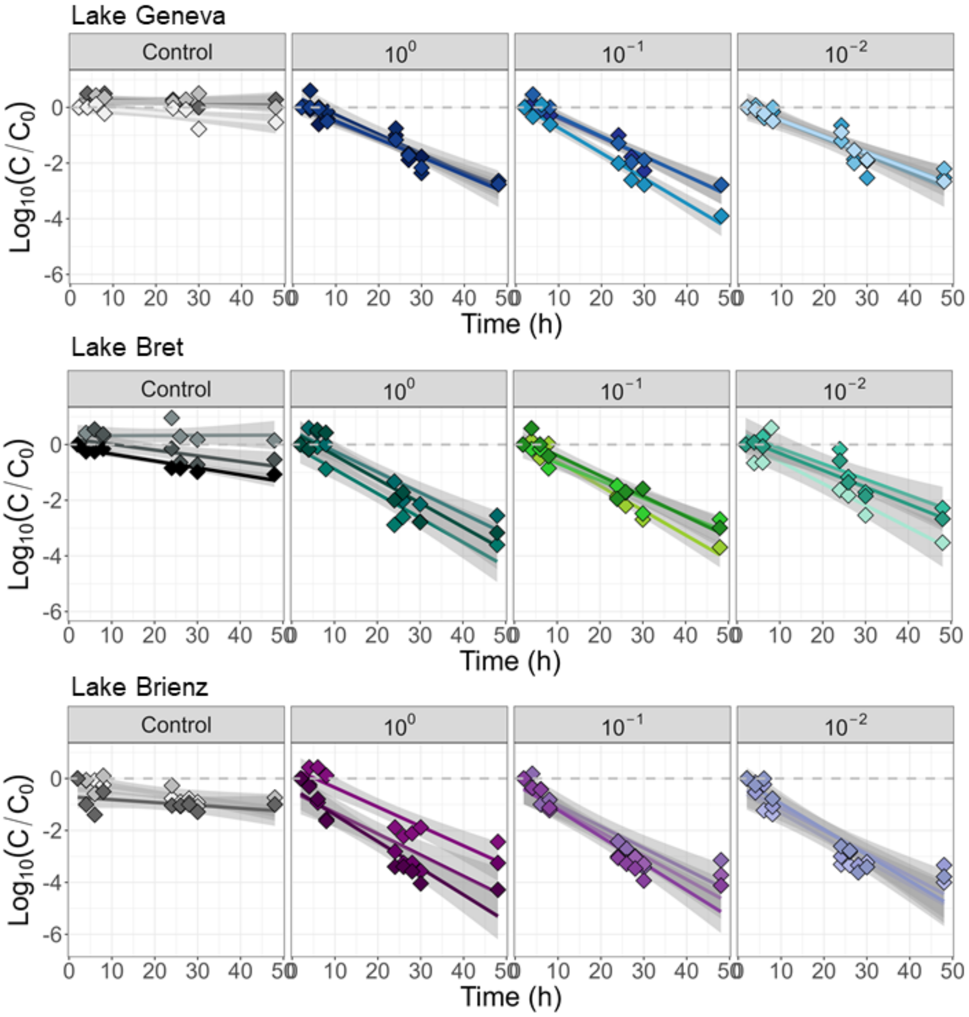

***Figure S5. Inactivation of Coxsackievirus B5 (CVB5) in lakewater chemostats.*** *After five days of growth, lakewater chemostats were spiked with a mixture of Coxsackievirus B5 (CVB5) and Human Adenovirus 2 (HAdV2). Samples were taken from the effluent to assess viral infectivity of CVB5 against Buffalo Green Monkey Kidney (BGMK) cells. Inactivation is shown as the loss of infectious virus concentration between the first sampling time and each time point ($C/C_0$). Each line represents a linear fit to the infectivity data for an individual chemostat and the shaded band the 95% confidence interval.*

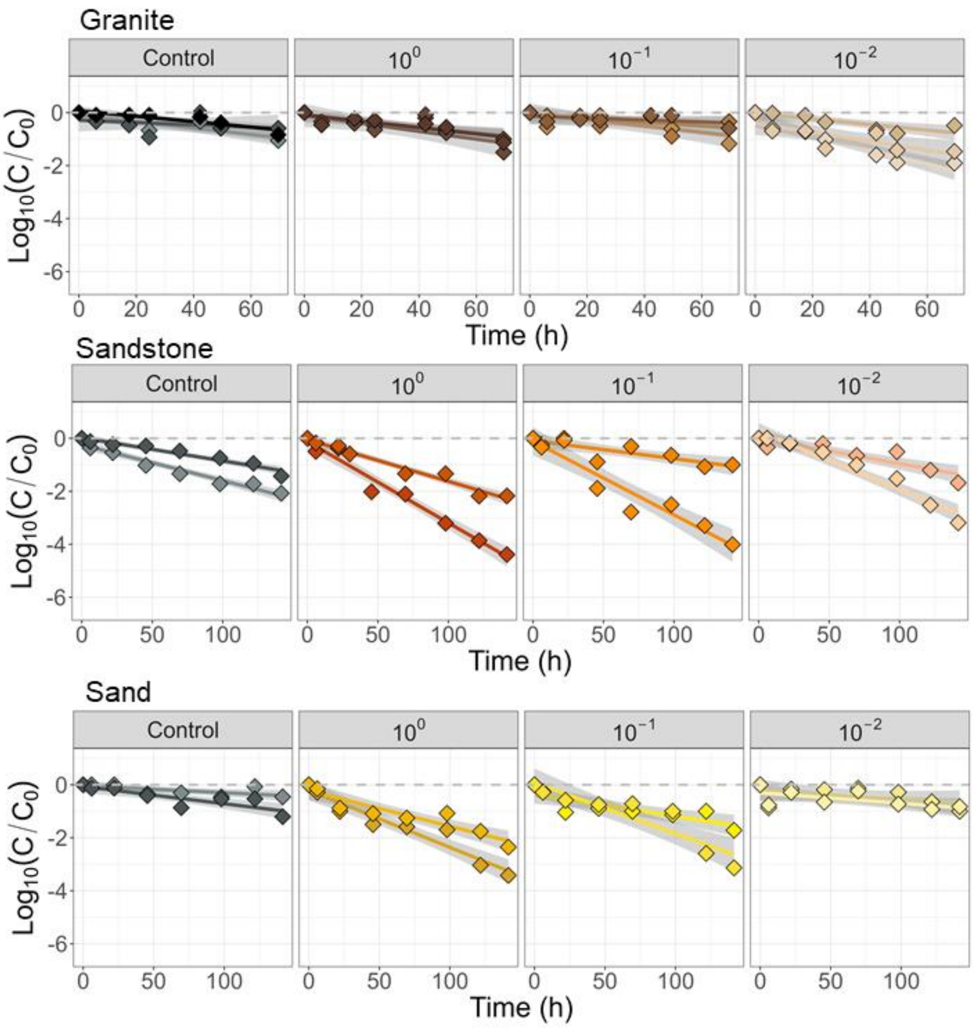

***Figure S6. Inactivation of Coxsackievirus B5 (CVB5) in groundwater chemostats.*** *After seven days of growth, groundwater chemostats were spiked with a mixture of Coxsackievirus B5 (CVB5) and Human Adenovirus 2 (HAdV2). Samples were taken from the effluent to assess viral infectivity of CVB5 against Buffalo Green Monkey Kidney (BGMK) cells. Inactivation is shown as the loss of infectious virus concentration between the first sampling time and each time point ($C/C_0$). Each line represents a linear fit to the infectivity data for an individual chemostat and the shaded band the 95% confidence interval.*

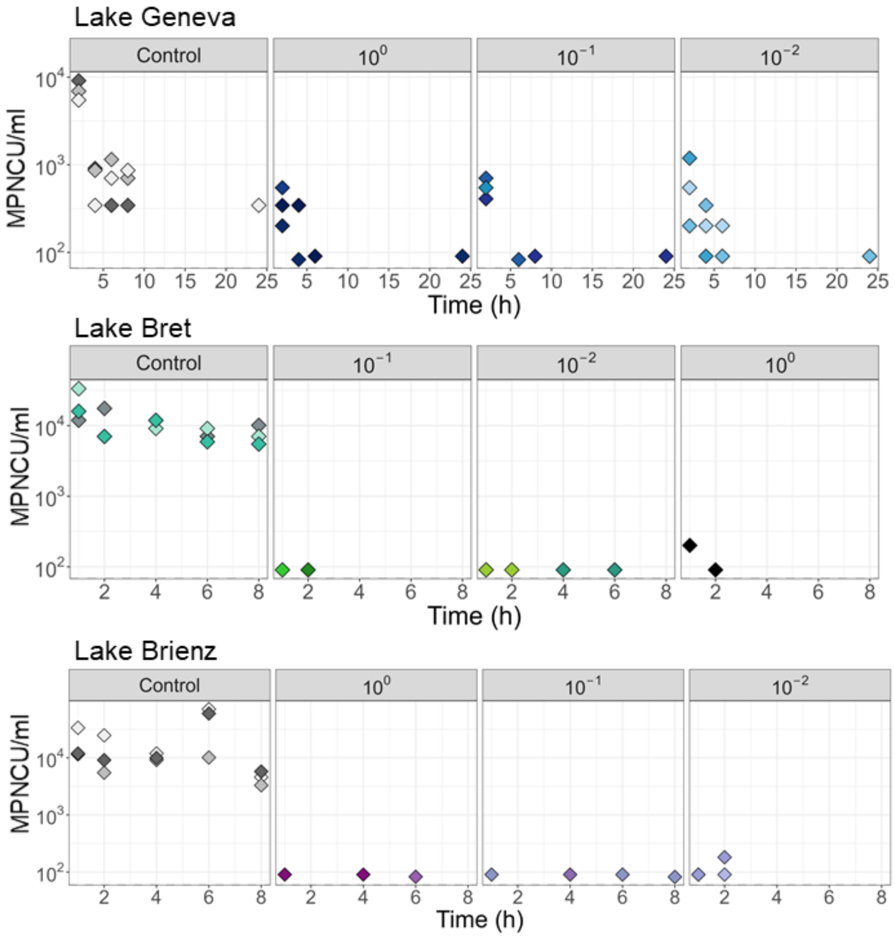

***Figure S7. Inactivation of Human Adenovirus 2 (HAdV2) in lakewater.*** *After five days of growth, groundwater chemostats were spiked with a mixture of Coxsackievirus B5 (CVB5) and Human Adenovirus 2 (HAdV2). Samples were taken from the effluent to assess viral infectivity of HAdV2 against rhabdomyosarcoma (RD) cells. Viral infectious cytopathic units (MPNCU/ml) over time were calculated using most probable number (MPN).*

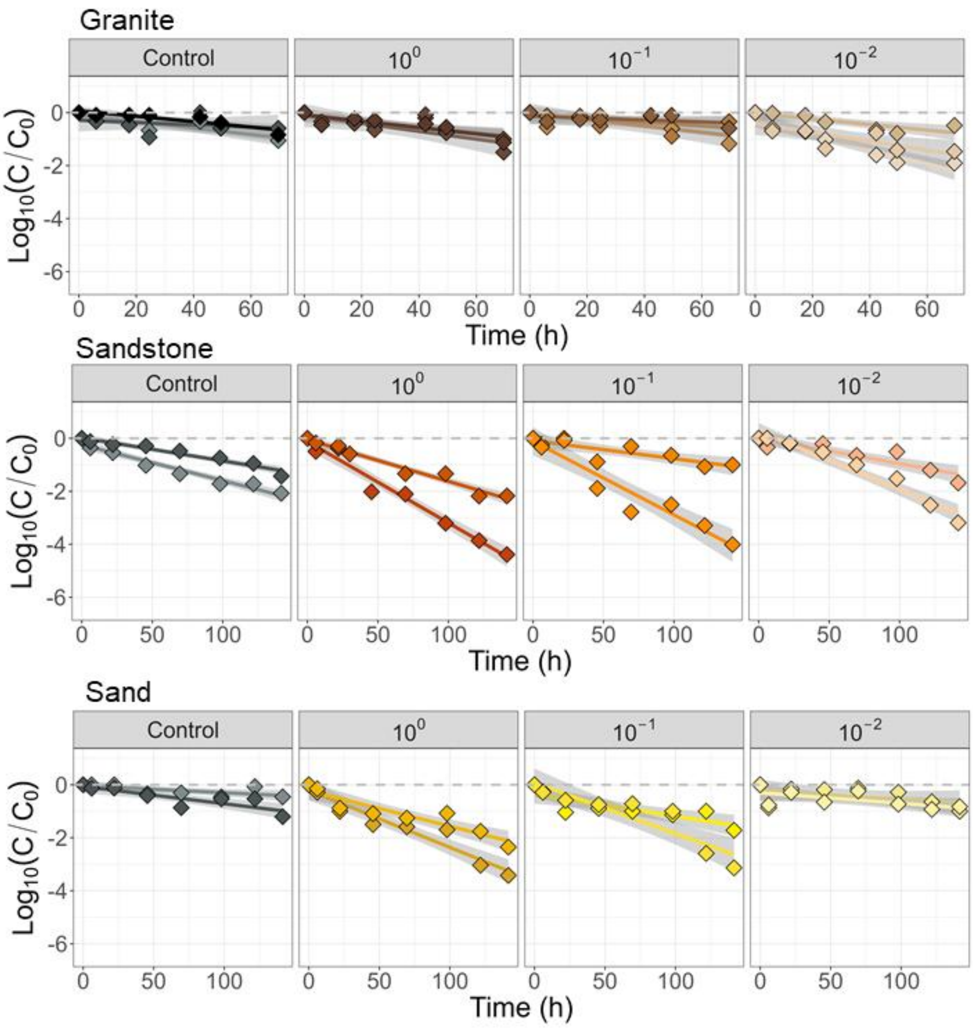

***Figure S8. Inactivation of Human Adenovirus 2 (HAdV2) in groundwater.*** *After seven days of growth, groundwater chemostats were spiked with a mixture of Coxsackievirus B5 (CVB5) and Human Adenovirus 2 (HAdV2). Samples were taken from the effluent to assess viral infectivity of HAdV2 against rhabdomyosarcoma (RD) cells. Inactivation is shown as the loss of infectious virus concentration between the first sampling time and each time point ($C/C_0$). Each line represents a linear fit to the infectivity data for an individual chemostat and the shaded band the 95% confidence interval.*

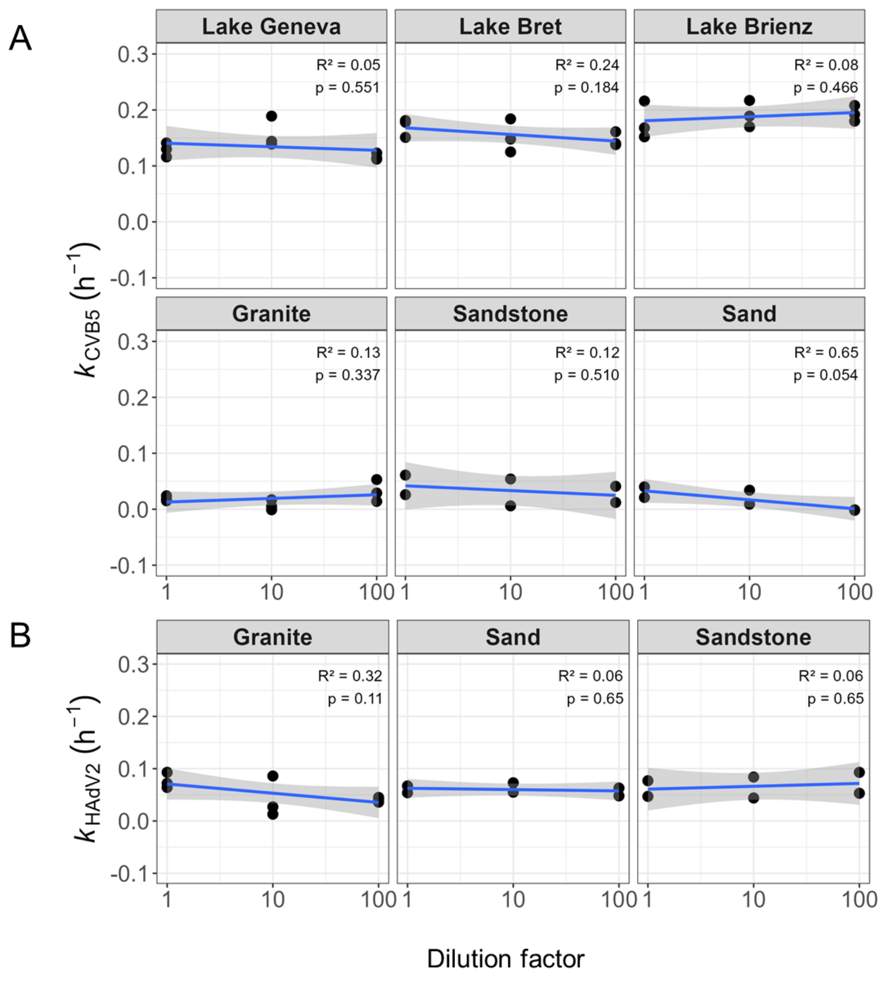

***Figure S9. Log-linear correlation of calculated inactivation rate constants and starting dilution factors****. $k_{CVB5}$* ***(A)*** *and $k_{HAdV2}$* ***(B)*** *were plotted as a function of the seeding dilution factor.*

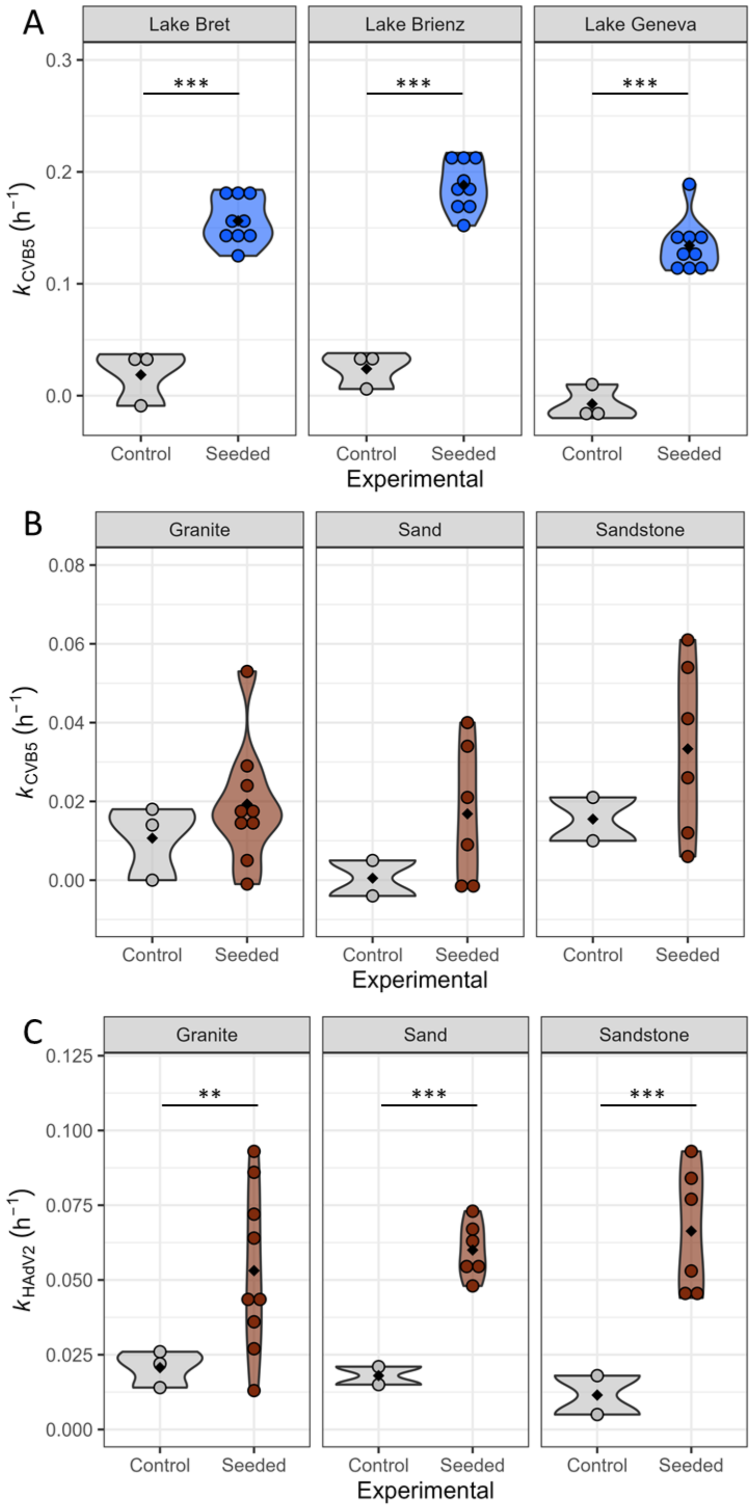

***Figure S10. Inactivation of unseeded controls and seeded chemostats.*** *Differences in the calculated inactivation rate constants between control and seeded chemostats were assessed using Welch's two-sample t-test. Comparison of calculated $k_{CVB5}$ in lakewater chemostats* ***(A)****, $k_{CVB5}$ in groundwater chemostats* ***(B)*** *and $k_{HAdV2}$ in groundwater chemostats* ***(C)****. Significance codes: 0 '***' 0.001 '**' 0.01 '*' 0.05.*

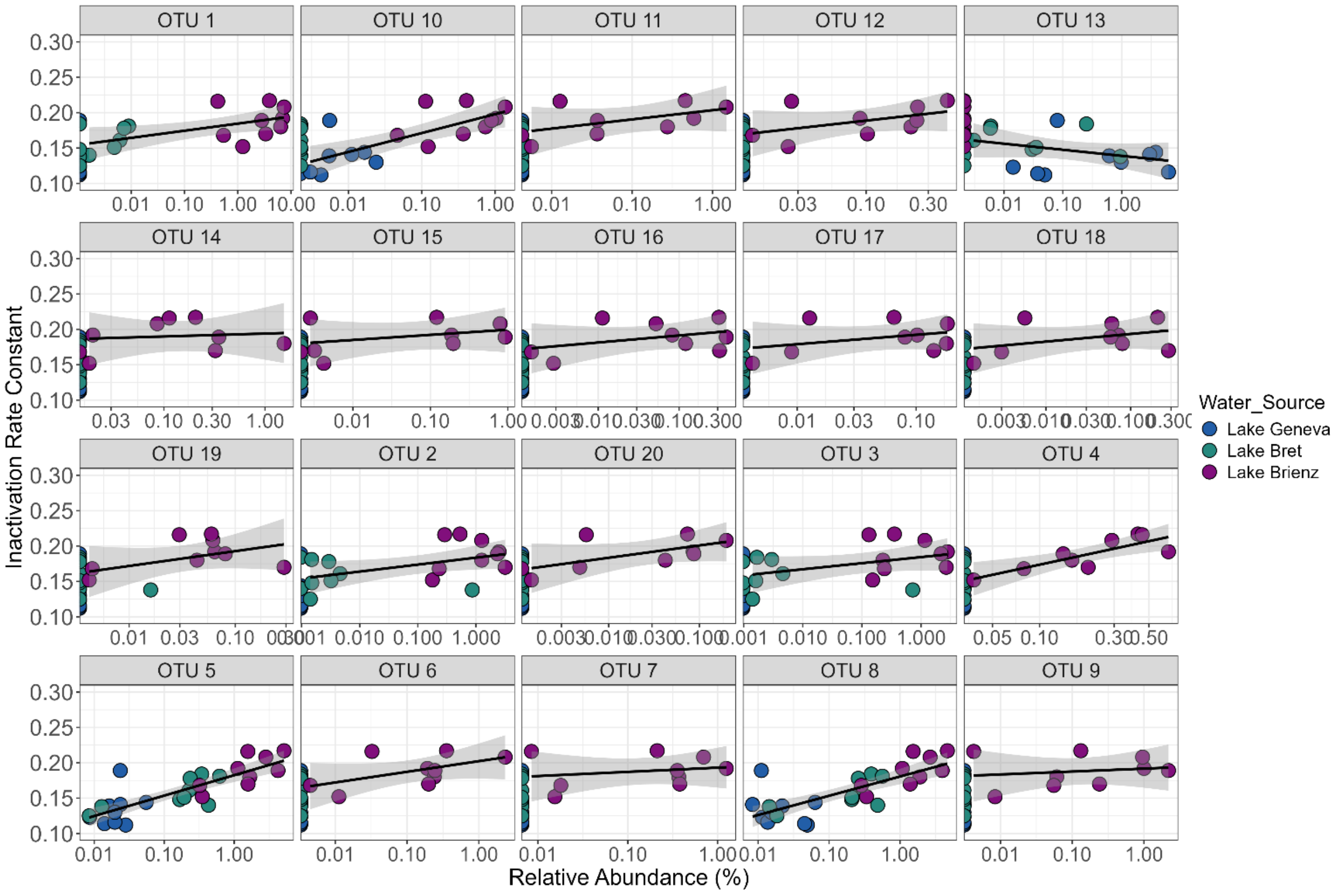

***Figure S11. Top 20 taxa with the largest estimated abundance changes as a function of $k_{CVB5}$ across lakewater chemostats.***

**Table S13. Taxonomy classification of top 20 taxa with the largest estimated abundance changes as a function of calculate CVB5 rate constants across lakewater chemostats.**

| OTU | beta | qval | direction | Family | Genus |
|---|---|---|---|---|---|
| OTU 1 | 2.495931 | 0.000531 | Positive | f__Comamonadaceae | g__ |
| OTU 2 | 1.90727 | 0.002924 | Positive | f__Burkholderiaceae | g__Limnobacter |
| OTU 3 | 1.802221 | 0.003526 | Positive | f__Burkholderiaceae | g__Limnobacter |
| OTU 4 | 1.767389 | 0.000628 | Positive | f__Comamonadaceae | g__ |
| OTU 5 | 1.713863 | 1.55E-05 | Positive | f__Comamonadaceae | g__Acidovorax |
| OTU 6 | 1.660727 | 0.00489 | Positive | f__Comamonadaceae | g__ |
| OTU 7 | 1.584342 | 0.008959 | Positive | f__Caulobacteraceae | g__Brevundimonas |
| OTU 8 | 1.571417 | 0.000335 | Positive | f__Comamonadaceae | g__Acidovorax |
| OTU 9 | 1.553082 | 0.018985 | Positive | f__Comamonadaceae | g__ |
| OTU 10 | 1.54904 | 0.006071 | Positive | f__Caulobacteraceae | g__Phenylobacterium |
| OTU 11 | 1.54073 | 0.01247 | Positive | f__Comamonadaceae | g__Paucibacter |
| OTU 12 | 1.488667 | 0.002924 | Positive | f__Caulobacteraceae | g__Brevundimonas |
| OTU 13 | -1.472 | 0.039334 | Negative | f__Rhodobacteraceae | g__Gemmobacter |
| OTU 14 | 1.470139 | 0.0022 | Positive | f__Comamonadaceae | g__Acidovorax |
| OTU 15 | 1.362972 | 0.028138 | Positive | f__Comamonadaceae | g__ |
| OTU 16 | 1.332027 | 0.006071 | Positive | f__Caulobacteraceae | g__Phenylobacterium |
| OTU 17 | 1.266579 | 0.003526 | Positive | f__Beijerinckiaceae | g__Bosea |
| OTU 18 | 1.194389 | 0.007549 | Positive | f__Comamonadaceae | g__ |
| OTU 19 | 1.137406 | 0.0022 | Positive | f__Burkholderiaceae | g__Limnobacter |

| | | | | | |
|---|---|---|---|---|---|
| OTU 20 | 1.136824 | 0.01247 | Positive | f__Comamonadaceae | g__ |

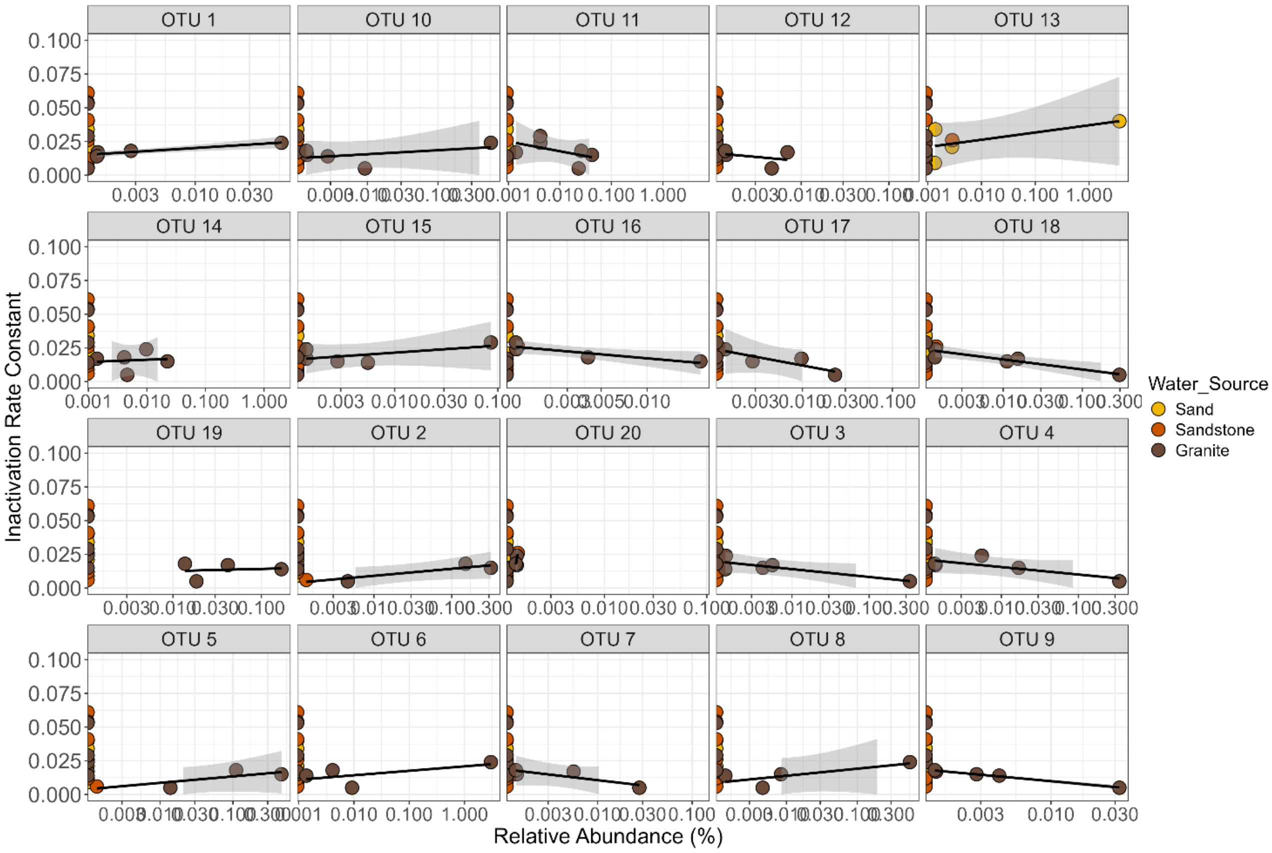

***Figure S12. Top 20 taxa with the largest estimated abundance changes as a function of $k_{CVB5}$ across groundwater chemostats.***

**Table S14. Taxonomy classification of top 20 taxa with the largest estimated abundance changes as a function of $k_{CVB5}$ across groundwater chemostats.**

| OTU | beta | qval | direction | Family | Genus |
|---|---|---|---|---|---|
| OTU 1 | 7.433032 | 0.001718 | Positive | f__ | g__ |
| OTU 2 | 5.335888 | 0.002494 | Positive | f__Sphingomonadaceae | g__ |
| OTU 3 | -4.74401 | 0.002494 | Negative | f__Comamonadaceae | g__Aquabacterium |
| OTU 4 | -4.31717 | 0.002494 | Negative | f__Oxalobacteraceae | g__Massilia |
| OTU 5 | 4.310342 | 0.002867 | Positive | f__Sphingomonadaceae | g__Sphingobium |
| OTU 6 | 4.153407 | 0.002796 | Positive | f__Comamonadaceae | g__Acidovorax |
| OTU 7 | -3.948 | 0.002494 | Negative | f__Comamonadaceae | g__Aquabacterium |
| OTU 8 | 3.808823 | 0.002867 | Positive | f__Methylophilaceae | g__Methylotenera |
| OTU 9 | -3.77998 | 0.002494 | Negative | f__Rhodocyclaceae | g__Dechloromonas |
| OTU 10 | 3.769467 | 0.002961 | Positive | f__Methylophilaceae | g__Methylotenera |
| OTU 11 | -3.62225 | 0.002217 | Negative | f__Comamonadaceae | g__Acidovorax |
| OTU 12 | -3.43679 | 0.004062 | Negative | f__Comamonadaceae | g__ |
| OTU 13 | 3.410059 | 0.006719 | Positive | f__Comamonadaceae | g__Hydrogenophaga |
| OTU 14 | -3.32138 | 0.003587 | Negative | f__Comamonadaceae | g__Acidovorax |
| OTU 15 | 3.234959 | 0.002838 | Positive | f__Comamonadaceae | g__ |
| OTU 16 | -3.23387 | 0.002636 | Negative | f__Sphingomonadaceae | g__Altererythrobacter |
| OTU 17 | -3.19986 | 0.002961 | Negative | f__Comamonadaceae | g__Aquabacterium |
| OTU 18 | -3.07602 | 0.001718 | Negative | f__Comamonadaceae | g__Aquabacterium |
| OTU 19 | 2.996758 | 0.003263 | Positive | f__Comamonadaceae | g__Aquabacterium |
| OTU 20 | -2.97754 | 0.002867 | Negative | f__Sphingomonadaceae | g__Sphingopyxis |

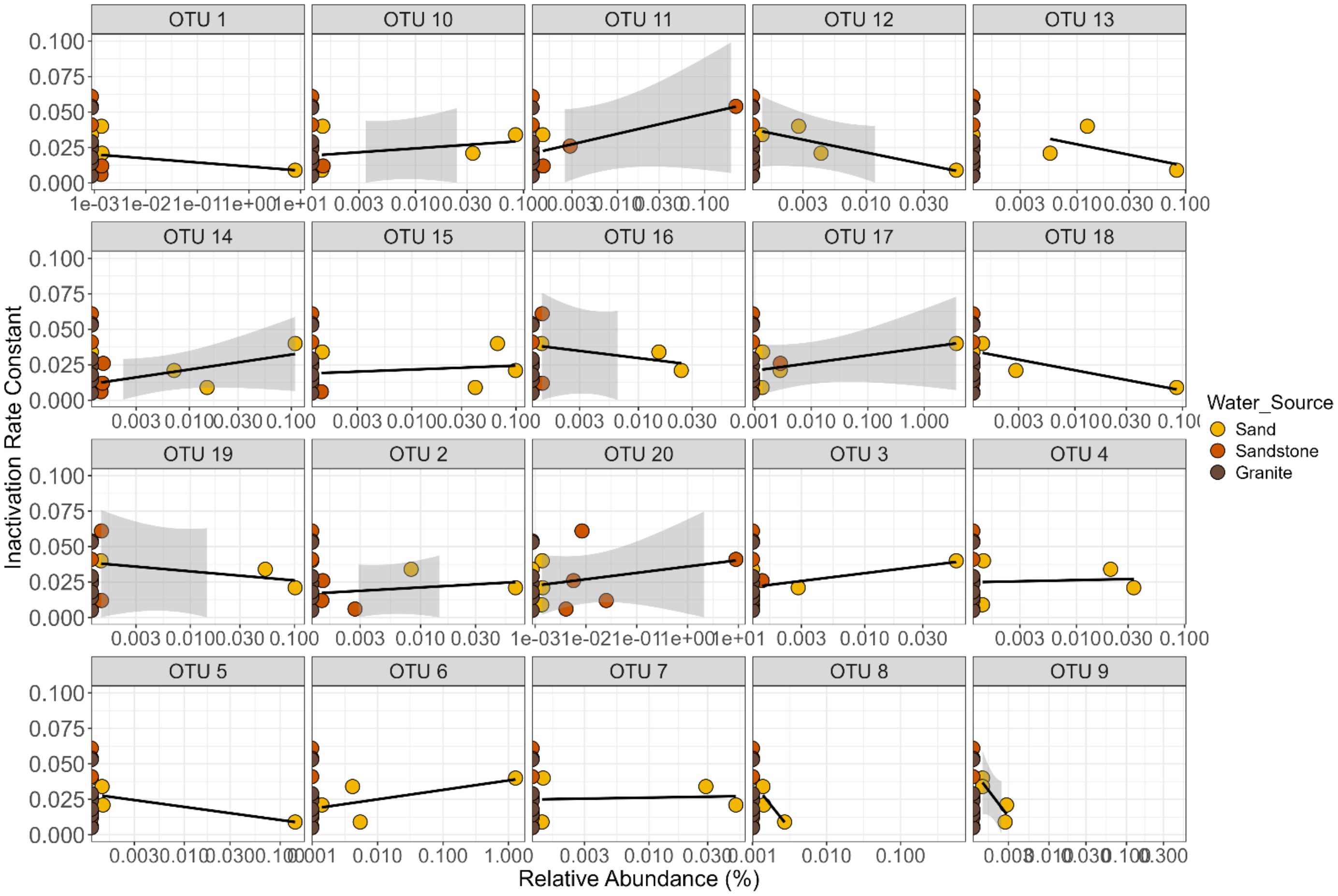

***Figure S13. Top 20 taxa with the largest estimated abundance changes as a function of $k_{HAdV2}$ across groundwater chemostats.***

**Table S15. Taxonomy classification of top 20 taxa with the largest estimated abundance changes as a function of $k_{HAdV2}$ across groundwater chemostats.**

| OTU | beta | qval | direction | Family | Genus |
|---|---|---|---|---|---|
| OTU 1 | 4.406641 | 0.001339 | Positive | f__Comamonadaceae | g__Aquabacterium |
| OTU 2 | 4.065662 | 0.001406 | Positive | f__Sphingomonadaceae | g__Sphingorhabdus |
| OTU 3 | 3.955298 | 0.002584 | Positive | f__Comamonadaceae | g__Aquabacterium |
| OTU 4 | -3.95375 | 0.002584 | Negative | f__Comamonadaceae | g__Aquabacterium |
| OTU 5 | 3.784529 | 0.003131 | Positive | f__Comamonadaceae | g__ |
| OTU 6 | 3.283982 | 0.002348 | Positive | f__Comamonadaceae | g__Aquabacterium |
| OTU 7 | -3.26906 | 0.001648 | Negative | f__Sphingomonadaceae | g__ |
| OTU 8 | -3.15422 | 0.004915 | Negative | f__Comamonadaceae | g__Aquabacterium |
| OTU 9 | -3.06531 | 0.002152 | Negative | f__Comamonadaceae | g__Aquabacterium |
| OTU 10 | -2.92113 | 0.001339 | Negative | f__Sphingomonadaceae | g__ |
| OTU 11 | 2.864835 | 0.002152 | Positive | f__Comamonadaceae | g__Acidovorax |
| OTU 12 | 2.787892 | 0.004258 | Positive | f__Comamonadaceae | g__Aquabacterium |
| OTU 13 | 2.760472 | 0.004258 | Positive | f__Comamonadaceae | g__Hydrogenophaga |
| OTU 14 | 2.754452 | 0.002152 | Positive | f__Comamonadaceae | g__Aquabacterium |
| OTU 15 | 2.539408 | 0.002152 | Positive | f__Comamonadaceae | g__Hydrogenophaga |
| OTU 16 | -2.53204 | 0.002321 | Negative | f__Comamonadaceae | g__ |
| OTU 17 | 2.478327 | 0.010054 | Positive | f__Comamonadaceae | g__Hydrogenophaga |

| | | | | | |
|---|---|---|---|---|---|
| OTU 18 | 2.47417 | 0.005443 | Positive | f__Comamonadaceae | g__ |
| OTU 19 | -2.44182 | 0.002321 | Negative | f__Sphingomonadaceae | g__ |
| OTU 20 | 2.407218 | 0.005627 | Positive | f__Comamonadaceae | g__Aquabacterium |
